# The illusion of diversity: sampling design drives conflicting estimates of soil bacterial richness

**DOI:** 10.64898/2026.07.03.736139

**Authors:** Maria Kostakou, Niklas Neisse, Kezia Goldmann, Antonis Chatzinotas, Stephanie D. Jurburg

## Abstract

Soil microbial diversity is shaped by the spatial scale at which communities are sampled, yet standard sampling practices often homogenize samples, obscuring fine-scale spatial structure and diversity patterns. To better understand how sampling effort, spatial extent, and physical homogenization influence plot-level microbial richness estimates, we sampled 57 forest and grassland sites across three regions in Germany using a 14-core cross-transect design, and performed 16S rRNA gene metabarcoding. We simulated sampling efforts of 1–14 cores and a range of spatial extents, and compared diversity estimates to those from physically homogenized composite samples. Plot-level richness increased continuously with sampling effort and spatial extent, with no evidence of saturation. However, when sequencing depth was held constant, sampling completeness declined with increasing sampling effort, meaning that more diversity is not captured. Composite samples substantially underestimated plot-level richness and altered apparent diversity relationships between ecosystems; individual cores identified forests as richer than grasslands, whereas homogenized samples suggested the opposite relationship. These results demonstrate that sampling effort, spatial extent, and homogenization fundamentally shape soil microbial diversity estimates. Homogenized composite samples cannot substitute for individual cores when the goal is to reliably quantify plot-level richness or compare diversity across ecosystems.

## Introduction

Soil microbial communities are exceptionally diverse and spatially heterogeneous (Fierer & Jackson 2006; Nunan *et al*. 2020), making them inherently prone to undersampling (Torsvik & Øvreås 2002), even when standard field and sequencing methods are applied. Macroecological studies have long shown that limited sampling systematically underestimates true richness, particularly in systems dominated by rare or patchily distributed organisms (Gotelli & Colwell 2001; Magurran 2021). In soils, the observed diversity of microbial communities increases steeply with sampling effort (Hermans *et al*. 2019; Joos *et al*. 2020), yet even intensive sampling protocols recover only a fraction of existing taxa, reflecting the enormous unseen diversity present in soils (Fierer & Jackson 2006; Delgado-Baquerizo *et al*. 2016). DNA extraction, amplification, and sequencing protocols further constrain the ability to fully characterize soil microbial communities, for example by setting upper limits on the amount of soil from which DNA can be extracted (Jurburg *et al*. 2021, 2022). Therefore, a major challenge is to design sampling strategies that adequately capture spatial variability and allow reliable estimation of microbial diversity.

Given existing limitations, mixing or homogenizing multiple soil cores from a single plot is a common practice aimed at increasing the representation of the plot-level microbial diversity while reducing sequencing costs and laboratory effort (Thompson *et al*. 2017; Jurburg *et al*. 2021). However, this practice obscures the plot-level spatial distribution of these communities. Diversity estimates directly depend on the spatial scale of sampling, which is in turn defined by the sample grain (i.e., the mass, volume, or area of samples taken; (Bach *et al*. 2018)), the spatial extent of a study, and the distance between samples (Dungan *et al*. 2002; Chase *et al*. 2018). In soils, especially when using composite samples, the grain is not defined by a single parameter, but rather by the number of cores homogenized (i.e., sampling effort) and the area over which the cores are taken (i.e., usually defined as a plot). When multiple soil cores are merged into a single composite sample, limits on DNA extraction protocols generally dictate that the same sample mass is taken per sample (usually <5 g of soil) regardless of sampling effort. Furthermore, sequencing depth limitations and the need for rarefaction (Schloss 2024) result in equal numbers of observations across samples regardless of the plot size or sampling effort. While rarefaction remains one of the most reliable ways to control for uneven sequencing effort (Weiss *et al*. 2017; Schloss 2024), it does not resolve biases introduced by differences in plot size or homogenization (Chase & Knight 2013; Chase *et al*. 2018). Larger plots, especially when sampled with greater effort, are therefore expected to detect higher diversity, consistent with the taxa–area relationship (Wiens 1989; Levin 1995; Drakare *et al*. 2006; Castle *et al*. 2019). Indeed, coverage-based approaches provide a framework for quantifying sampling completeness and show that substantially greater effort is theoretically required to fully characterize highly diverse soil microbial assemblages (Chao & Jost 2012).

Importantly, as the taxa-area relationship is non-linear (Wiens 1989; Levin 1995), homogenization introduces ambiguity by removing fine-scale spatial information that structures microbial communities (Ettema 2002; Castle *et al*. 2019). This compromises diversity estimates derived from composite samples taken over different plot areas or sampling efforts, and complicates direct comparisons between composite and individual soil cores (Castle *et al*. 2019). This limitation is particularly important as long-term and global efforts for monitoring soil biodiversity are established (Fischer *et al*. 2010; Guerra *et al*. 2021; Parnell *et al*. 2025; Singh *et al*. 2025).

Although experimental work has demonstrated that homogenizing soil decreases local richness and reduces between-sample variance (West & Whitman 2022), its impact as a standard field sampling practice on the detection of soil microbial diversity remains unexplored, particularly across different sampling efforts and plot sizes. To assess how these methodological choices shape microbial diversity estimates in soils, we took 855 soil samples in 57 sites across three regions of Germany, spanning a gradient of land-use intensities in both grasslands and forests. Soils in these long-term research sites (Biodiversity Exploratories; Fischer *et al*. 2010) have been sampled regularly for over a decade using a fixed-distance orthogonal transect design with 14 soil cores (seven per transect) evenly distributed along each transect. At each site, we took composite soil samples as well as individual cores for each point in the transect and performed 16S rRNA gene metabarcoding for composite and individual samples, following standard protocols. Simulating homogenization *in silico* and comparing the resulting soil bacterial communities to those derived from physical composite samples and individual soil cores, we aimed to quantify how plot size and sampling effort influence the detection of soil microbial diversity and determine whether the process of DNA extraction and rarefaction introduces biases in diversity detection.

## Methods

### Study sites and sample collection

Soil samples were collected from the Biodiversity Exploratories (Fischer *et al*. 2010), a long-term research platform to study biodiversity and ecosystem processes across environmental and land-use gradients in Germany. The three Exploratories (Schwäbische Alb, Hainich-Dün, and Schorfheide-Chorin) each contain 100 sites (50 grassland, 50 forest) representing a range of management intensities and environmental conditions. Within this framework, a subset of 57 sites is designated as Very Intensive Plots (VIPs), representing core study sites with extensive ecological and environmental background data. This subset includes nine forest and nine grassland sites per Exploratory, with three additional forest sites in Hainich-Dün.

Soil sampling in these VIPs followed a standardized spatial protocol to capture fine-scale within-site heterogeneity. Each forest plot measures 100 × 100 m, while each grassland plot measures 50 × 50 m. Soil sampling was conducted in May 2023. At each site, 14 individual soil cores were collected along two orthogonal transects (north-south and west-east) using a 5 cm diameter soil auger from the 0–10 cm soil layer. In forests, samples were spaced every 6 m along 40 m transects; in grasslands, samples were spaced every 3 meters along 20 m transects. Approximately 5 g of soil was reserved with a spoon per core for individual sample analysis by proportionally subsampling soil from the entire 0–10 cm core depth to obtain a representative sample. True composite samples were created by physically homogenizing the rest of the 14 individual cores with a 2 mm sieve into a single bulk sample per plot prior to DNA extraction. Individual core samples were not sieved. All samples were stored at −20 °C until DNA extraction.

### DNA extraction, library preparation, sequencing, and bioinformatics

DNA was extracted from 0.3 g of soil per sample using the Nucleospin Soil Kit (Macherey-Nagel, Düren, Germany), following the manufacturer’s protocol. DNA concentrations were quantified with a *Qubit 4 Fluorometer using the dsDNA High Sensitivity (HS) Assay Kit* (Invitrogen, Thermo Fisher Scientific). All DNA extracts were stored at –20 °C until further processing. Amplicon libraries for bacteria and archaea were prepared using a two-step dual-indexed PCR targeting the V4 region of the 16S rRNA gene, following the standard Earth Microbiome Project protocols (Thompson *et al*. 2017). The first PCR employed the primers 16S_Illu_515F (5′-TCGTCGGCAGCGTCAGATGTGTATAAGAGACAGGTGYCAGCMGCCGCGGTAA-3′) and 16S_Illu_806R (5′-GTCTCGTGGGCTCGGAGATGTGTATAAGAGACAGGGACTACNVGGGTWTCTAAT-3′), which included Illumina overhang adapter sequences. Sample-specific indices were added in a second PCR using the Nextera XT Index Kit (Illumina). Negative controls were included in all PCRs to monitor contamination. PCR products were purified using AMPure XP magnetic beads (Beckman Coulter), quantified again with the Qubit system, and pooled. Final libraries were sequenced on an Illumina MiSeq platform using 2 × 250 bp paired-end reads (MiSeq Reagent Kit v3 600 cycles, Illumina Inc., San Diego, CA, USA) at the Department of Applied Microbial Ecology, Helmholtz Centre for Environmental Research, Leipzig, Germany.

All bioinformatics processing was done with the *DADA2* pipeline (Callahan *et al*. 2016) in R version 4.4.1 (R Core Team 2024). Paired-end reads were filtered and trimmed using the *FilterAndTrim* function with truncLen = c(160, 160), trimLeft = c(10, 10), trimRight = c(10, 10), maxN = 0, maxEE = c(3, 3), and truncQ = 2, with all other parameters kept at default values. Error rates were learned independently for forward and reverse reads, and ASVs were inferred per sample. Paired reads were merged, and chimeras were removed using consensus-based detection. Taxonomy was assigned using the SILVA v138.1 database (Quast *et al*. 2013) with the DADA2 classifier. Processed sequences were analyzed using the *phyloseq* (McMurdie & Holmes 2013) and *vegan* (Jari Oksanen 2007) packages. All samples were rarefied to 8,000 reads prior to analysis; samples below this threshold were excluded. Consequently, the number of plots available for analysis differed between individual and composite sample metrics at maximum sampling effort (N = 14), with 22 forest and 22 grassland plots retained for individual core analyses and 25 forest and 24 grassland plots for composite sample analyses.

### Diversity assessments

For each plot, we randomly subsampled, without replacement, 1–14 individual cores to obtain a gradient of *in silico* composite sampling efforts. Subsampling was repeated 100 times per sampling effort, per plot to account for stochastic variation in core selection. Diversity was quantified for each subsample by summing ASV counts across selected individual soil cores to calculate *in silico* composite ASV richness, which served as an estimate of plot-level richness. Summing samples to calculate ASV richness linearly increased the number of sequences per composite sample with sampling effort. Therefore, to disentangle the effect of increased sequencing depth from increased sampling effort (Gotelli & Colwell 2001; Weiss *et al*. 2017; Schloss 2024), all *in silico* composite samples were additionally standardized to a fixed depth of 8,000 reads, equivalent to the sequencing depth of a single sample before richness estimation. Unlike its conventional use to control for uneven sequencing effort among independent samples (Schloss 2024), in this case, standardization served to simulate the fixed sequencing depth applied to composite samples regardless of the number of cores pooled, testing the consequence of this common laboratory practice.

In addition to richness, we quantified sampling completeness with Good’s coverage, which measures the proportion of the total community represented by richness and is less sensitive to sampling intensity than raw richness alone (Good 1953; Chao & Jost 2012) for individual and *in silico* composite samples using the *coverage()* estimator implemented in *betaC* (Engel *et al*. 2021), both before and after sequencing-depth standardization. Individual cores (x = 1) were considered as sample-level richness, while the true composite samples were taken as estimates of plot-level richness. Within-plot heterogeneity was calculated as the ratio of plot-level to sample-level richness for each subsample. For each sampling effort and plot, diversity metrics were averaged across the 100 subsampling replicates before subsequent analyses.

To quantify information loss associated with sequencing-depth standardization, the ratio of standardized to unstandardized plot-level richness was calculated at three representative sampling efforts spanning the gradient (x = 3, 6, and 12 cores), providing a measure of the proportion of richness retained after standardization. Differences in the variance of plot-level richness between ecosystem types were assessed using Levene’s test (Fox *et al*. 2024).

To further examine the effect of spatial configuration between samples on diversity estimates derived from *in silico* composite samples, we calculated the sum of all pairwise Euclidean distances (in meters) among the selected samples, providing a quantitative measure of the spatial spread of sampling within a plot at a given sampling effort.

### Statistical analyses

To examine how plot-level richness responded to increasing sampling effort, we fitted generalized linear mixed models (GLMMs; Gamma error distribution, log link) using the *glmmTMB* package (Brooks *et al*. 2017). Sampling effort (number of samples), ecosystem type (forest vs. grassland), and their interaction were included as fixed effects, while plot identity was modeled as a random intercept. Equivalent models were fitted to richness estimates obtained after sequencing-depth standardization. To assess whether plot-level richness showed evidence of saturation with increasing sampling effort, we compared linear, log-linear, square root, and asymptotic models of the richness-sampling effort relationship using AIC, with ΔAIC > 2 considered meaningful support for the better-fitting model (Guthery *et al*. 2003). Model performance was evaluated using conditional and marginal R² values (*performance* package; (Lüdecke *et al*. 2021). The significance of predictors was tested using Type III Wald χ² tests (Fox *et al*. 2024), and model assumptions were assessed using simulated residuals (*DHARMa;* Hartig et al., 2024). Predicted means, 95% confidence intervals, and ecosystem-specific slopes along log-transformed sampling effort were estimated using the *emmeans* package (*emmeans()* and *emtrends()* functions; Lenth 2024). Contrasts between slopes were used to test whether richness and coverage accumulation differed between forests and grasslands. For models using log transformed predictors, slopes were back-transformed to express the multiplicative change associated with a doubling of sampling effort or spatial extent. Descriptive statistics (mean ± SD) were further calculated for minimum (x = 1) and maximum (x = 14) sampling efforts per ecosystem. Where distributions were skewed, median and interquartile range (IQR) are reported instead of mean ± SD. Median ratios of grassland to forest richness were calculated as effect sizes for ecosystem comparisons at overlapping sampling efforts.

To assess how sampling effort influenced sampling completeness (calculated as plot-level coverage), we fitted GLMMs (Beta error, logit link) for observed and standardized plot-level coverage, with the same model structure as described above. Sampling effort (number of samples), ecosystem type, and their interaction were included as fixed effects, and plot identity as a random intercept. Model fit, assumption checks, and significance testing followed the same procedures. Predicted means and 95% confidence intervals were used for visualization, and descriptive statistics were computed for the lowest and highest sampling efforts in each ecosystem.

To explore how species richness varied with spatial extent, we fitted GLMMs with a Gamma error distribution and log link. Spatial extent was summarized as the mean extent per plot across subsampling replicates before analysis. Log-transformed sampling extent, ecosystem type, and their interaction were included as fixed effects, and plot identity was modeled as a random intercept. We conducted model diagnostics, inference, and performance evaluation as described previously. Predicted means and 95% confidence intervals were calculated to compare the magnitude and rate of increase in richness across spatial scales. To assess whether plot-level richness showed evidence of saturation with increasing sampling extent, we compared four models of the relationship using AIC as above. To assess how spatial extent influenced plot-level coverage, we fitted equivalent GLMMs (Beta error, logit link) with log-transformed spatial extent, ecosystem type, and their interaction as fixed effects, and plot identity as a random intercept. Model fit, diagnostics, and inference followed the same procedures as described above.

To quantify the scaling of richness from sample to plot level, we modeled plot-level richness as a function of log-transformed sample-level richness, log-transformed spatial extent, ecosystem type, and their interactions as fixed effects, with plot identity as a random intercept (GLMM; Gamma error, log link) across three spatial extent classes (0–75 m, 100–300 m, and 600–1,000 m). Estimated marginal means were extracted for the representative extents to summarize ecosystem-specific patterns, and ratios of plot-level to sample-level richness were derived from model predictions to quantify differences in spatial scaling between forests and grasslands.

Finally, to test for ecosystem differences in plot-level richness while controlling for spatial extent, richness values were averaged to plot-level means, and analyses were restricted to the subset of spatial extents shared by both ecosystem types at each sampling effort. Because the spatial extent covered by a given sampling effort differs between forest and grassland plots due to their different spatial configurations, overlap was only present for sampling efforts of 2–7 samples, where both ecosystem types were fully represented and spatial extents remained within a comparable range. Wilcoxon rank-sum tests were performed separately for each sampling effort, with Holm-adjusted p-values to account for multiple comparisons, and median ratios of grassland to forest richness were reported as effect sizes.

## Results

### Sampling effort effects on plot-level richness and coverage

To assess how sampling effort influences the microbial diversity observed at each site, we modeled plot-level richness as a function of sampling effort (i.e., the number of samples randomly taken per site). Plot-level richness increased with sampling effort in both grassland and forest ecosystems (Figure 1a). For individual samples derived from a single soil sample, we observed 1,411 ± 911 ASVs in forests, on average, compared to 675 ± 515 ASVs in grasslands. With the highest sampling effort (14 samples), we detected 10,546 ± 4,267 ASVs in forests and 6,341 ± 2,432 ASVs in grasslands (Figure 1a). Model comparison strongly favored a log-linear relationship between sampling effort and plot-level richness over an asymptotic model (ΔAIC = 975), providing no evidence of richness saturation within the range of sampling efforts examined (1–14 samples). After standardization, richness increased to 4,178 ± 718 ASVs in forests and 3,314 ± 548 ASVs in grasslands (Figure 1b; Figure S3). For standardized richness, the log-linear and asymptotic models were statistically indistinguishable (ΔAIC = 0.84).

**Figure 1.**
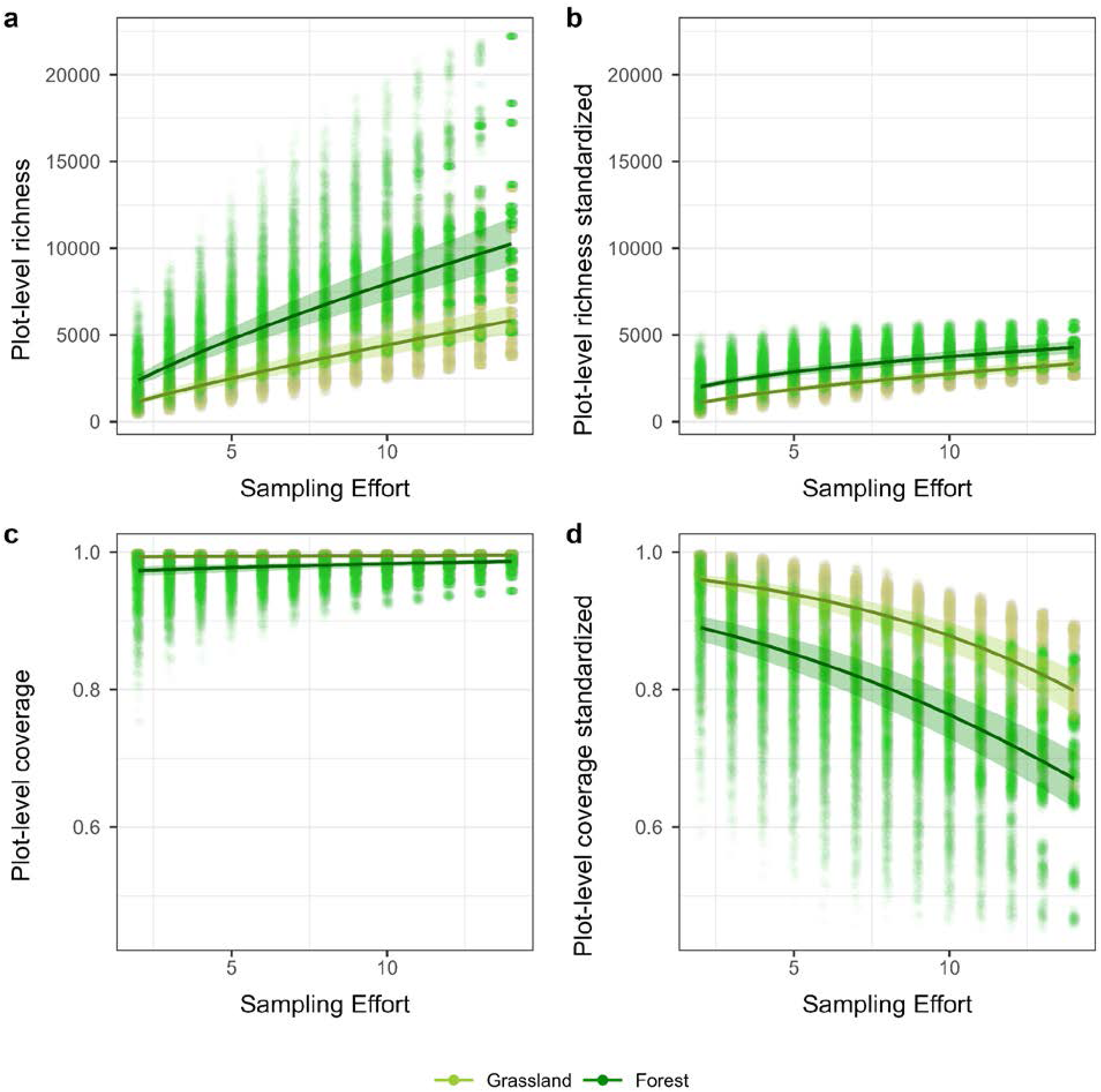
Plot-level richness and plot-level coverage as a function of sampling effort (2–14 samples) in forest and grassland soils. **(a)** Plot-level richness, **(b)** plot-level species richness standardized to a fixed sequencing depth of 8,000 reads, **(c)** plot-level coverage, and **(d)** plot-level coverage of the depth-standardized community. Points represent individual replicate subsamples per plot; lines show model-fitted relationships from generalized linear mixed models, and shaded areas indicate 95% confidence intervals (CI).

Plot-level richness was strongly influenced by sampling effort, ecosystem type, and their interaction, all of which were highly significant (Gamma mixed-effects models, log link; *p* < 0.001; R²_conditional_ = 0.931; R²_marginal_ = 0.627; Table S1). Among these effects, sampling effort explained the largest proportion of variation in richness, while ecosystem type and its interaction with sampling effort accounted for additional, but smaller, differences between forest and grassland plots. Doubling the number of cores increased plot-level richness by a factor of 1.70 in forests (95% CI: 1.703 – 1.709) and 1.79 in grasslands (95% CI: 1.780 – 1.800), with forests reaching higher absolute richness across all sampling efforts.

We observed a similar overall pattern when richness was calculated after standardizing sequencing depth (Gamma mixed-effects models, log link; *p* < 0.001; R²_conditional_ = 0.890; R²_marginal_ = 0.579). Doubling the number of cores increased standardized plot-level richness by a factor of 1.34 in forests (95% CI: 1.338–1.345) and 1.53 in grasslands (95% CI: 1.519–1.531). Importantly, fewer ASVs were detected after standardization, with richness at maximum sampling effort reduced from 10,546 ± 4,267 to 4,178 ± 718 ASVs in forests (a 60% reduction) and from 6,341 ± 2,432 to 3,314 ± 548 ASVs in grasslands (a 48% reduction), indicating that standardization disproportionately reduced detectable richness in forests relative to grasslands.

To assess the effect of finite sequencing depth on diversity estimates across increasing field sampling effort, we calculated plot-level coverage. Plot-level coverage with a single sample averaged 0.948 ± 0.061 in forests and 0.987 ± 0.025 in grasslands, and increased slightly with additional sampling (Figure 1c). At the highest effort (14 samples), plot-level coverage increased to 0.984 ± 0.013 in forests and 0.994 ± 0.006 in grasslands. Plot-level coverage increased with sampling effort, with significant effects of sample size, ecosystem type, and their interaction (beta GLMM with logit link; all p < 0.001; Table S2), indicating ecosystem-specific differences in coverage accumulation, with grasslands reaching higher coverage at lower sampling effort than forests (Figure 1c; Figure S4). Plot-level coverage increased by a factor of 1.061 in forests (95% CI: 1.060 – 1.062) and by 1.029 in grasslands (95% CI: 1.027 – 1.031) with each additional sample, with a significant difference in slopes between ecosystems (z = −33.1, p < 0.0001) (Figure 1c).

Notably, after standardizing sequencing depth across sampling effort, plot-level coverage declined with increasing sampling effort, in contrast to the increase observed for non-standardized richness (Figure 1d). After sequencing-depth standardization, coverage decreased with increasing sampling effort, reaching 0.699 ± 0.100 in forests and 0.808 ± 0.067 in grasslands with 14 samples (Figure S5). This decline was significantly steeper in forest than in grasslands, supported by a highly significant interaction between sample size and ecosystem type (p < 0.0001). Each additional sample decreased standardized plot-level coverage by a factor of 0.891 in forests (95% CI: 0.891–0.892) and 0.861 in grasslands (95% CI: 0.860–0.862), with a significantly steeper decline in grasslands (z = −60.2, p < 0.0001) (Figure 1d).

### Composite sampling and information loss

To assess information loss from physical sample homogenization, we compared the richness in true composite samples with richness estimates derived from individual cores, including mean sample-level richness, *in silico* plot-level richness, and sequencing-depth-standardized plot-level richness (Figure 2a-c; Figure S6). Consistent with the patterns described above, forests exhibited higher richness than grasslands across all individual-core-based metrics (Holm-adjusted p < 0.001). At the sample level, we detected 1,377 ± 595 ASVs per sample in forests (n = 22 plots) compared to 731 ± 339 ASVs in grasslands (n = 22 plots; Figure 2a), based on plot-level means at maximum sampling effort (x = 14). Notably, this relationship was reversed in true composite samples (Figure 2d), where grasslands showed higher richness (1,109 ± 840 ASVs) than forests (655 ± 317 ASVs; Wilcoxon rank-sum test: W = 197, Holm-adjusted p = 0.040). True composite samples recovered a median of 6.2% (IQR: 6.1%) in forests and 16.2% (IQR: 22.5%) in grasslands of the richness detected on the *in silico* composite samples of the same 14 cores (Figure 3). The recovery ratios differed significantly between ecosystems (Wilcoxon rank-sum test: W = 136, p = 0.001).

**Figure 2.**
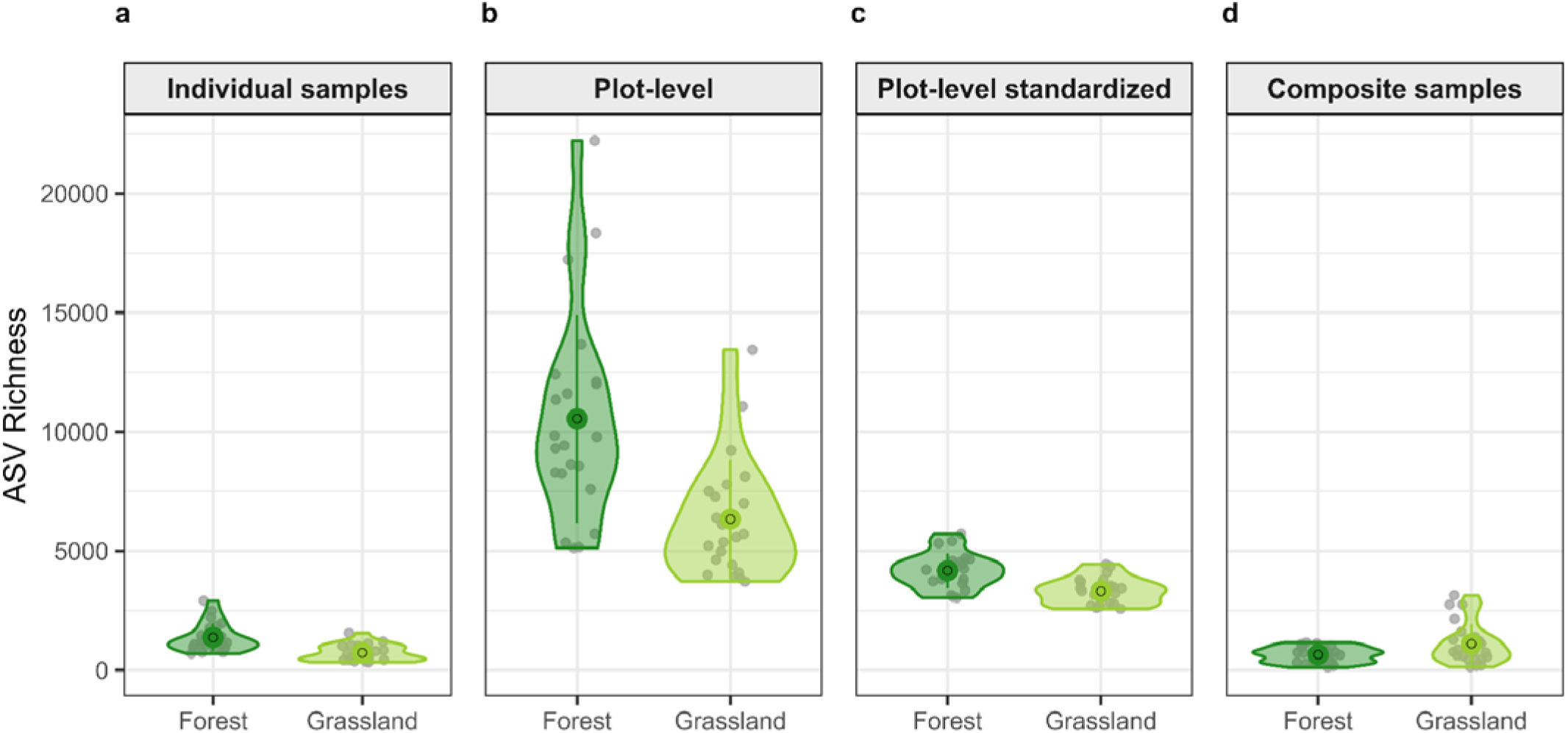
ASV richness at maximum sampling effort (N = 14 cores) across four metrics in forest and grassland soils. (a) Mean sample-level richness per plot, averaged across 14 individual cores. (b) Plot-level richness from the *in silico* composite of 14 cores. (c) Plot-level richness standardized to a fixed sequencing depth of 8,000 reads. For panels a–c, values represent plot-level means averaged across 100 random subsampling replicates. (d) ASV richness from one physically homogenized composite sample per plot. Violins show the distribution across plots; points and error bars show mean ± 1 SD across plots. Grey points represent individual plot values.

**Figure 3.**
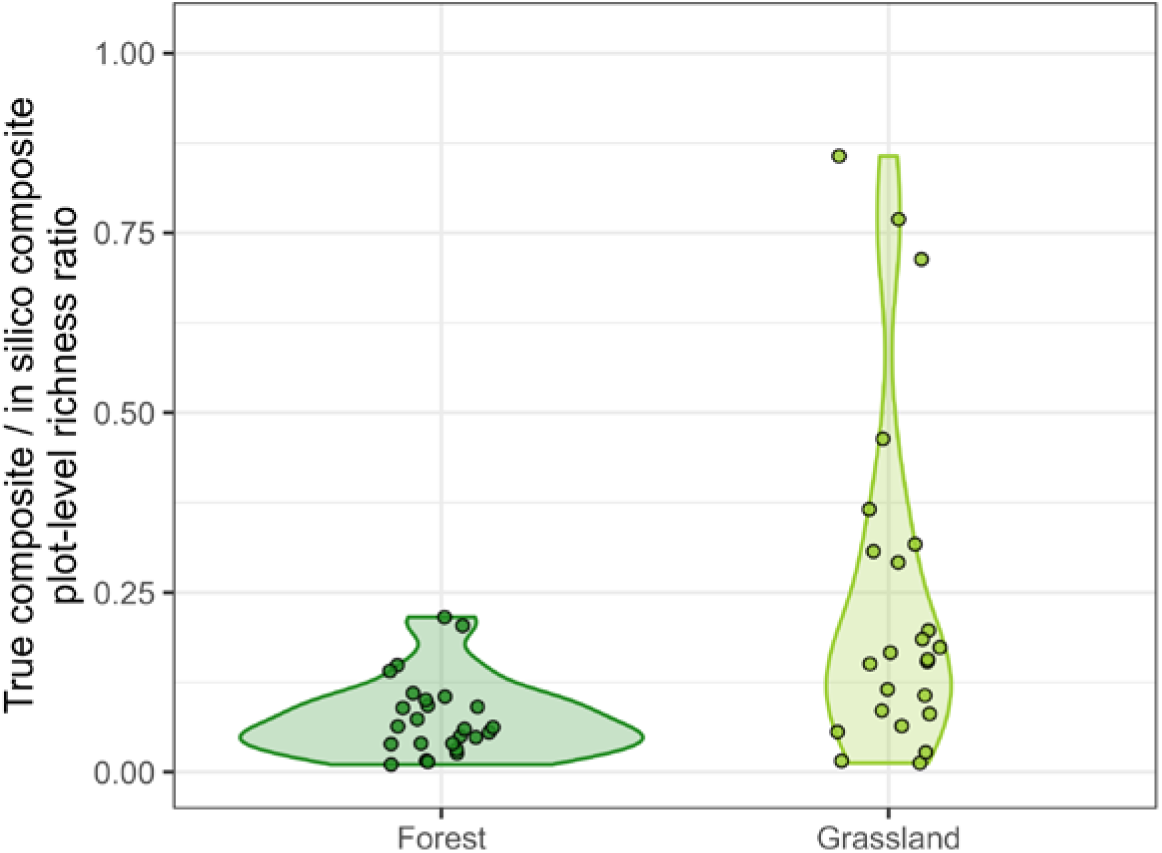
Recovery of plot-level ASV richness by true composite samples relative to *in silico* composites of the same 14 cores, across forest and grassland soils. Points represent composite plots; a ratio of 1 indicates complete capture of plot-level diversity by the composite sample.

We further quantified information loss associated with sequencing depth standardization across sampling efforts (Figure 4). Across all sampling efforts and both ecosystems, standardized richness estimates captured less diversity than unstandardized richness estimates, with this difference increasing with sampling effort. In forests, standardized composite samples created with 3, 6, and 12 samples recovered only 74% (95% CI: 71–77%), 58% (55–62%), and 45% (41–48%) of the diversity recovered with unstandardized composite samples, respectively. Similar patterns were found in grasslands, where standardized composite samples created with 3, 6, and 12 samples recovered 86% (84–89%), 74% (70–78%), and 60% (56–64%) of the diversity recovered with unstandardized composite samples, respectively. Differences in recovery between forests and grasslands were significant at all sampling efforts (Wilcoxon rank-sum test, all Holm-adjusted p < 0.001). The proportion of richness retained after standardization declined significantly with sampling effort in both ecosystems (Spearman’s ρ = −0.820 in forests and ρ = −0.799 in grasslands, both p < 0.0001; Figure 4), indicating that information loss from sequencing-depth standardization increased with the number of samples included in the composite sample. Although standardization reduced absolute richness values in both ecosystems, forests were consistently richer than grasslands, and this gap widened with sampling effort (Holm-adjusted p < 0.001; Figure S1, Figure S3).

**Figure 4.**
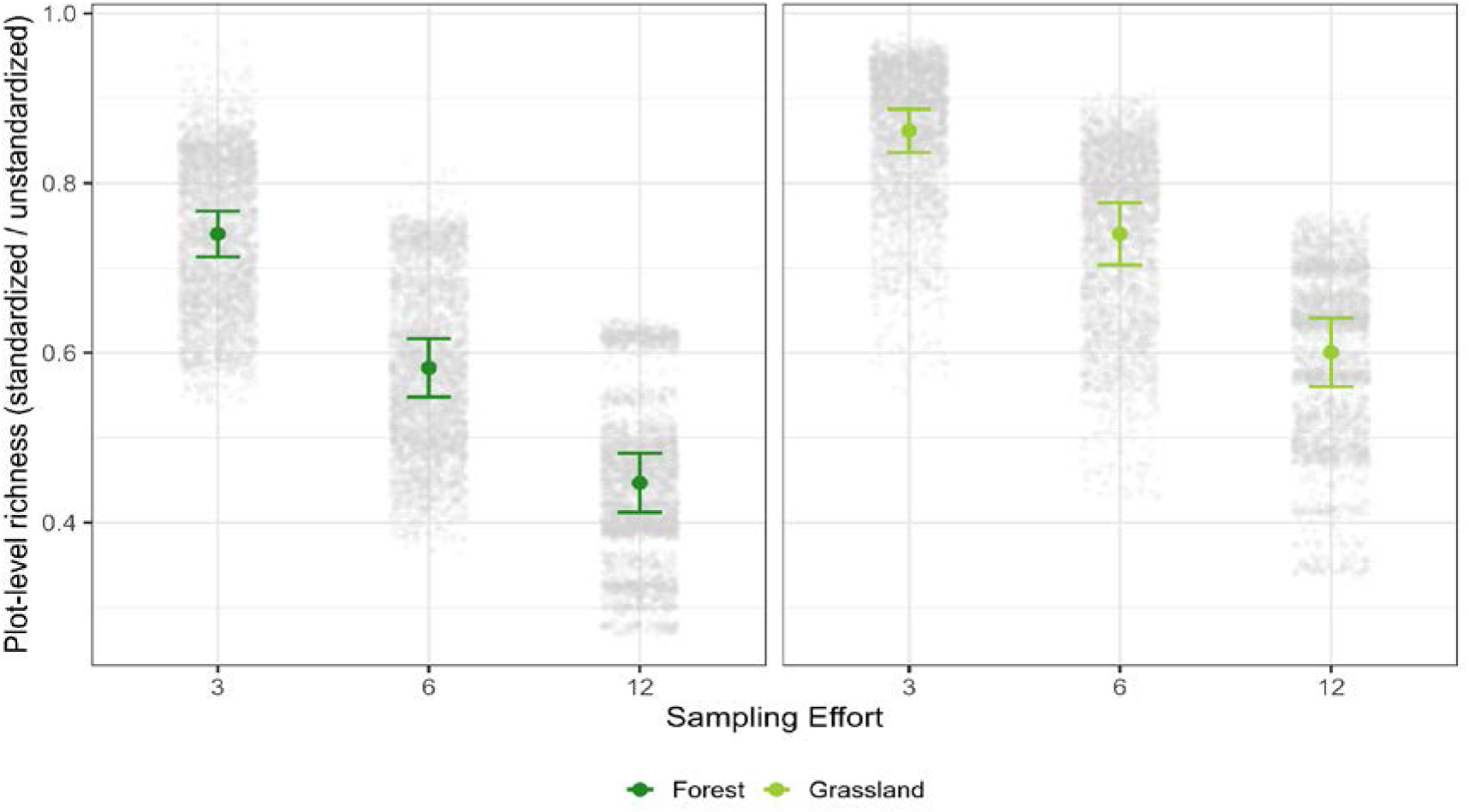
Ratio of standardized to unstandardized plot-level richness as a function of sampling effort. Gray points show all observations with replicates; colored points mark the mean ratio per sample size with ±95% CI across plots, shown separately for forest and grassland.

Within-plot heterogeneity, expressed as the ratio of plot-level to sample-level richness, increased with sampling effort in both ecosystems. However, it diverged between the two ecosystems with increasing sampling effort, reaching higher values in grasslands than in forests (forest: 2.5× to 7.7×; grassland: 2.6× to 8.7× across x = 3 to 12; Figure S7).

### Spatial scaling on plot-level richness and coverage

To assess how spatial extent influences plot-level richness and coverage, we modeled plot-level richness as a function of the spatial extent. Plot-level richness increased with spatial extent in both ecosystems, reaching higher absolute values in forests, but with a steeper rate of increase in grasslands (Figure 5a; z = 37.9, p < 0.0001). The effects of plot-level richness, spatial extent, ecosystem type, and their interaction were statistically significant (all p < 0.001; R²_conditional_ = 0.928; R²_marginal_ = 0.622; Table S3). Doubling the spatial extent increased plot-level richness by a factor of 1.24 in forests (95% CI: 1.240–1.242) and 1.27 in grasslands (95% CI: 1.264–1.267). Model comparison strongly supported a log-linear relationship between spatial extent and plot-level richness compared to alternative model forms (ΔAIC = 6,214.47; Table S6), and this held after sequencing-depth standardization (ΔAIC > 29,619.90; Table S6), indicating no evidence of richness saturation across the range of spatial extents considered. Standardized plot-level richness showed a similar but more gradual richness accumulation (Figure 5b; all p < 0.001; R²_conditional_ = 0.885; R²_marginal_ = 0.590). Doubling the spatial extent increased standardized plot-level richness by a factor of 1.12 in forests (95% CI: 1.118–1.120) and 1.18 in grasslands (95% CI: 1.175–1.177; z = 89.6, p < 0.0001).

**Figure 5.**
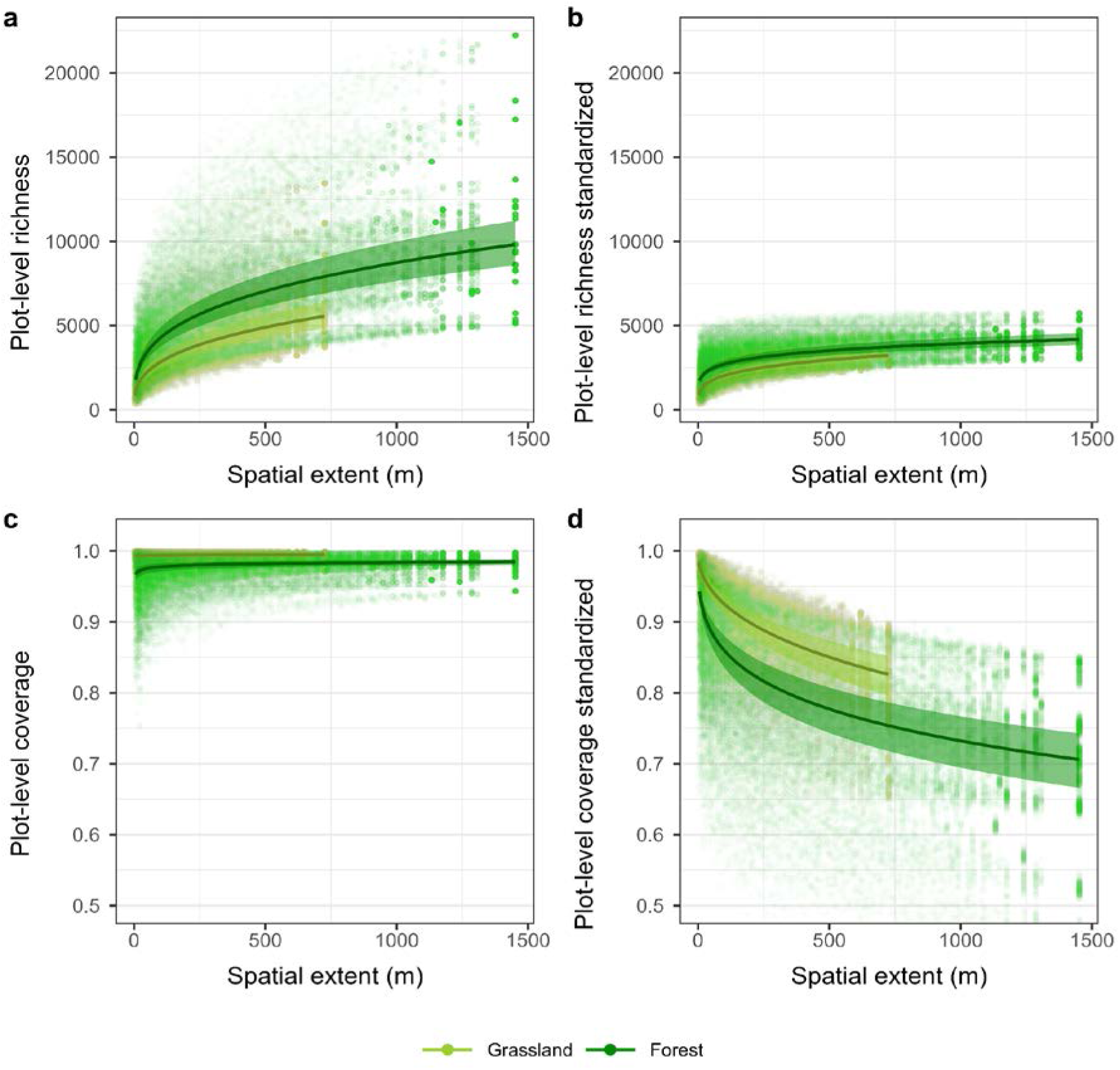
Plot-level richness and coverage as a function of spatial extent in forest and grassland soils. (a) Plot-level richness, (b) standardized plot-level richness (rarefied to 8,000 reads per composite sample), (c) plot-level coverage, and (d) standardized plot-level coverage. Points represent individual replicate subsamples per plot; lines show model-fitted relationships from generalized linear mixed models, and shaded areas indicate 95% confidence intervals (CI).

Sample coverage increased with spatial extent in both ecosystems, with steeper gains in forests (Figure 5c). Extent, ecosystem type, and their interaction had statistically significant effects on plot-level coverage (beta GLMM with logit link; all p < 0.001; R²_conditional_ = 1.000; R²_marginal_ = 0.428; Table S4). Doubling the spatial extent increased plot-level coverage by a factor of 1.10 in forests (95% CI: 1.103–1.106) and 1.05 in grasslands (95% CI: 1.050–1.056; *z* = −28.585, p < 0.0001). After standardizing sequencing depth, spatial extent, ecosystem type, and their interaction remained highly significant predictors of plot-level coverage (all p < 0.001; R²_conditional_ = 0.998; R²_marginal_ = 0.659). In contrast to richness, standardized plot-level coverage declined with increasing spatial extent. Specifically, doubling spatial extent resulted in proportional reductions in coverage to 0.784 in forests (95% CI: 0.783–0.785) and 0.715 in grasslands (95% CI: 0.714–0.716; *z* = −75.7, *p* < 0.0001) (Figure 5d).

To ensure comparable sampling effort between ecosystems, analyses were restricted to sampling efforts where both ecosystem types were fully represented (x = 2–7; Figure S8, Figure S9). In these sampling efforts, forest and grassland plots occupy different but partially overlapping ranges of spatial extent, as the fixed spacing difference between ecosystems (6 m vs 3 m) means forests always cover larger extents than grasslands for the same number of cores. Within this range, forest plots consistently exhibited higher microbial plot-level richness than grassland plots across all sampling efforts (Wilcoxon rank-sum tests, Holm-adjusted p < 0.001), with forests showing approximately twice the richness of grasslands (median grassland to forest ratio = 0.47–0.57). Although plot-level richness increased with sampling effort in both ecosystems, the absolute increase was greater in forests, reflecting a steeper accumulation of richness with spatial extent.

In addition, variance in plot-level richness was significantly greater in forests than in grasslands across all overlapping sampling efforts (Levene’s test, p < 0.05), indicating greater spatial heterogeneity among forest plots. Beyond x = 7, forest and grassland plots no longer co-occur within the same range of spatial extents (Figure S8), precluding direct ecosystem comparisons at broader sampling efforts.

### Scaling of plot-level richness with sample-level richness

To examine the relationship between sample-level and plot-level richness across spatial extents, we modeled their relationship across three extent classes (0–75 m, 100–300 m, and 600–1000 m). While previously we compared absolute richness between ecosystems at equivalent sampling efforts, here we focus on how richness scales from the sample to the plot level across increasing spatial extents, noting that forest and grassland plots reach equivalent spatial extents at different sampling efforts due to the fixed spacing difference between ecosystems. Across all spatial extents, plot-level richness scaled positively with sample-level richness in both ecosystem types (GLMM, Gamma error, log link; p < 0.001; Figure 6; Figure S10; Table S5). In both forest and grasslands, the slopes between sample- and plot-level richness decreased with spatial extent, from 1.09 and 1.15 at 0–75 m, to 0.98 and 0.96 at 100–300 m, and to 0.95 and 0.82 at 600–1000 m. This indicates that the proportional gain in plot-level richness per unit increase in sample-level richness diminished at broader spatial grains. Within the overlapping extent range, grassland plots showed steeper scaling than forests at the finest spatial extent (slopes: grassland = 1.15, forest = 1.09; difference = -0.057, p = 0.0012), while no significant ecosystem difference was detected at intermediate extents (100–300 m; difference = 0.020, p = 0.11), and forests showed steeper scaling at the broadest extent (600–1000 m; difference = 0.123, p < 0.001).

**Figure 6.**
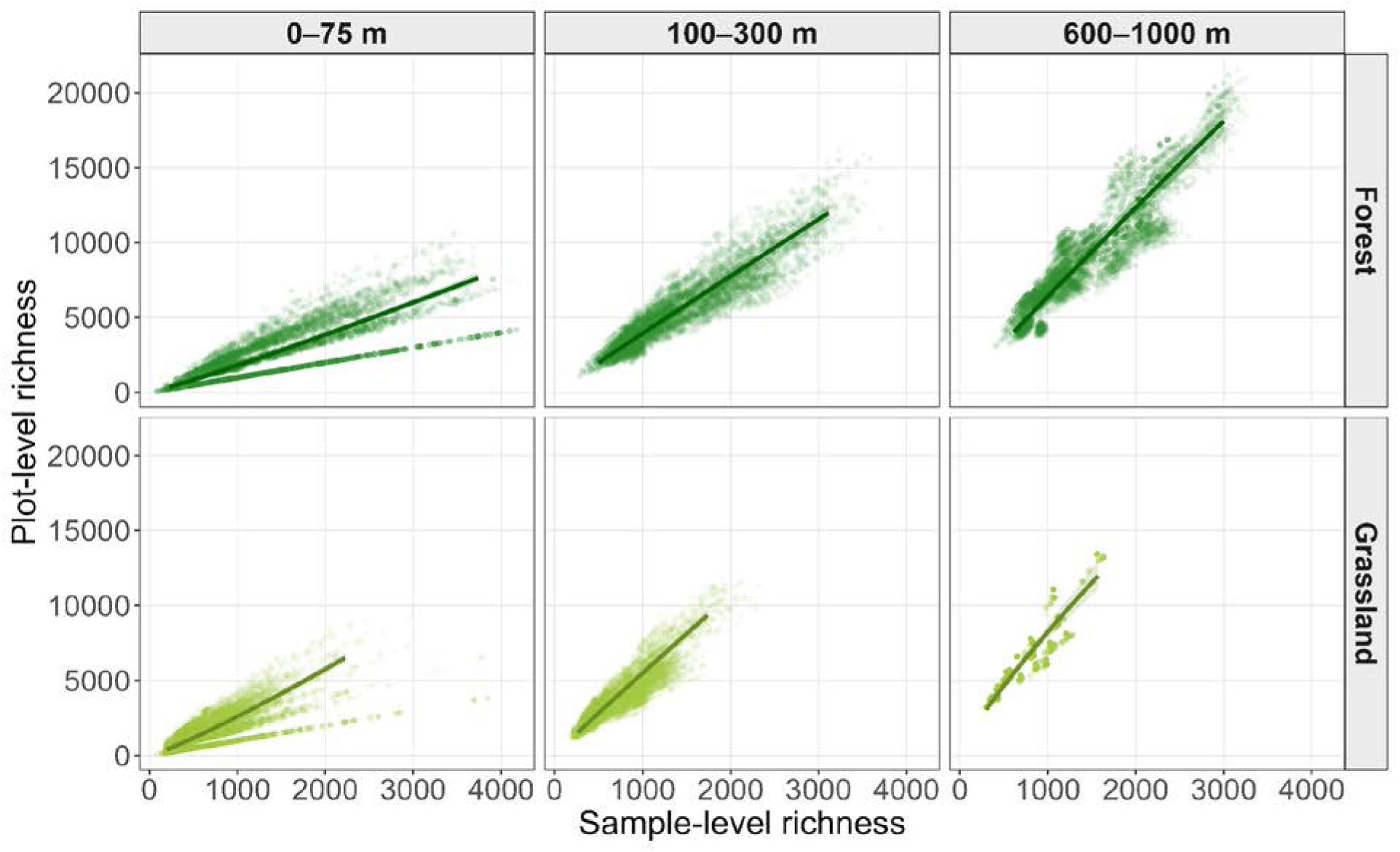
Relationship between sample-level and plot-level ASV richness across three spatial extent classes in forest and grassland soils. Points represent individual replicate subsamples per plot; lines and shaded bands show GLMM-fitted relationships with 95% confidence intervals (Gamma error, log link; plot as random intercept)

## Discussion

In ecology, sampling is rarely sufficient to detect all, or even most species present in an assemblage, particularly in assemblages dominated by large numbers of rare taxa (Gotelli & Colwell 2001; Magurran & McGill 2011) soil microbial communities, which at fine spatial scales are highly diverse and spatially heterogeneous, resulting in steep species–accumulation relationships and strong compositional turnover over short distances (Ettema 2002; Fierer & Jackson 2006; Baker *et al*. 2009). Τhis is not even accounting for additional methodological hurdles, such as the 0.5 g limit of most soil DNA extraction protocols, which restricts the amount of soil from which DNA can be extracted regardless of sampling effort. Given these difficulties, homogenizing multiple soil cores across a defined area has become standard in soil microbial ecology, with the implicit assumption that mixing more cores better captures the microbial diversity of a larger area (Robertson *et al*. 1999; Lawrence *et al*. 2020). Indeed, some studies have shown that the detection of soil microbiota increases with both spatial sampling effort and sequencing depth (Sogin *et al*. 2006; Lennon & Jones 2011; Shade *et al*. 2014; Jousset *et al*. 2017). Yet, comprehensive empirical tests of how sampling effort, spatial extent, and physical homogenization cumulatively affect soil microbial diversity estimates are lacking. We conducted an extensive field sampling across regions and ecosystems in Germany, simulated homogenization *in silico,* and compared diversity estimates to those obtained from physically homogenized samples to explore how well this process captures microbial diversity through metabarcoding. Consistent with ecological theory, we found that plot-level soil microbial richness is highly sensitive to how communities are sampled, with sampling effort, spatial extent, homogenization, and standardization each fundamentally altering diversity estimates, and in some cases, ecological relationships between plots. We discuss each of these findings in turn below.

Across all soils, plot-level richness increased continuously with each additional soil core (before sequencing-depth standardization), partly because each new core added more sequences to the *in silico* composites. However, imposing a sequencing depth standardization reversed this pattern: plot-level coverage declined as sampling effort increased, indicating that a fixed sequencing depth becomes less sufficient to capture the increasing number of low-abundance taxa introduced by additional cores. This was further supported by model comparison, where log-linear and asymptotic models were statistically indistinguishable for standardized richness (ΔAIC = 0.84), consistent with rarefaction to a fixed sequencing depth imposing an artificial ceiling on detectable richness. As homogenization is typically used to increase the representativity of a sample without increasing sequencing costs, enforcing this standardization allowed us to examine the effect of increasing field sampling effort independently of sequencing depth. This standardization is not simply an analytical choice; it represents a controlled *in silico* simulation of what occurs during physical compositing. When multiple soil cores are homogenized and then sequenced at a depth calibrated for a single sample, the diversity gained by adding more cores is systematically reduced. Since composite samples are typically processed using standard single-sample sequencing depths, this *in silico* approach provides a direct comparison with the most common practice. Our analysis demonstrates that diversity gains arising from sampling additional, spatially distinct soil cores are not captured by sequencing the resulting composite sample. This is likely due to methodological limitations associated with fixed sequencing depth (Chase & Knight 2013; Chase *et al*. 2018) and a fixed soil mass from which DNA is extracted, regardless of the mass of the homogenized sample.

Compared to the *in silico* composite samples, true composite samples substantially underestimated plot-level diversity, but this effect varied between ecosystems, leading to changes in the relationships between them. Specifically, the diversity of forest soils was underestimated to a greater extent than that of grassland soils, resulting in higher diversity in grasslands than forests in true composite samples, opposite to the pattern found across all three individual core-based metrics (Figure 2). This reversal is supported by the markedly lower recovery of diversity in true forest composite samples: forests recovered a median of only 6.2% of *in silico* plot-level richness compared to 16.2% in grasslands. Forest soils are characterized by high small-scale spatial heterogeneity (Baldrian 2017), suggesting that homogenization merges more compositionally distinct communities than in grasslands. Combined with the larger spatial extents covered by true forest composite samples at x = 14, more distinct communities are incorporated into the same fixed extraction mass, exacerbating the dilution of rare taxa. Physical homogenization reduces the detection of richness and spatial turnover by artificially inflating dispersal and reducing the spatial independence of microsites (Ranjard *et al*. 2013; Castle *et al*. 2019; West & Whitman 2022). Crucially, as composite sample mass increases, DNA extraction mass remains the same due to practical laboratory limitations, collapsing fine-scale spatial variability of multiple cores into a single, spatially unresolved observation (Wiens 1989; Levin 1995). It should also be noted that true composite samples were sieved through a 2 mm sieve prior to DNA extraction, whereas individual samples were processed without sieving. Sieving may have introduced additional changes in community composition beyond those attributable to physical homogenization alone, as it disrupts soil micro-aggregates that harbor distinct microbial communities (Bach *et al*. 2018). The observed differences in diversity estimates between individual and composite samples may therefore reflect the combined effects of homogenization and sieving rather than homogenization alone. Our study further supports the notion that physical homogenization obscures the spatial structure that characterizes soil microbial communities locally (Ettema 2002; Vos *et al*. 2013; Raynaud & Nunan 2014; Fierer 2017), and that these effects are particularly important in ecosystems with higher small-scale heterogeneity like forest soils (Baldrian 2017; Fierer 2017). Furthermore, we find that homogenization can mask ecologically meaningful diversity contrasts between ecosystems which is the aim of many long-term global monitoring programs (Fischer *et al*. 2010; Guerra *et al*. 2021; Parnell *et al*. 2025; Singh *et al*. 2025).

The diversity in a plot characterized by multiple cores reflects sample-level richness (i.e., our individual cores) and compositional turnover (Whittaker 1972; Drakare *et al*. 2006). We found that core sets encompassing greater spatial extents captured higher diversity across ecosystems, as predicted by the distance-decay relationship (Nekola & White 1999). Because the sampling geometry was identical across plots, these differences between grasslands and forests reflect ecosystem-specific differences in spatial turnover rather than artifacts of sampling design. Forests exhibited consistently higher plot-level richness than grasslands across spatial extents, while grasslands showed steeper richness accumulation per unit increase in spatial extent, suggesting higher spatial turnover in grasslands and greater overall heterogeneity in forests. Restricting comparisons to overlapping spatial extents shared by both ecosystem types confirmed a higher plot-level richness in forests than grasslands across all comparable sampling efforts. Crucially, we found no indication of saturation with increasing spatial extent, even after sequencing-depth standardization, highlighting the high spatial turnover of soil microbiota within the meter scale. This has direct implications for interpreting true composite samples: at the scale relevant to microorganisms, a single soil core already represents a complete, spatially discrete community, and distances between cores that appear small by macroecological standards are vast dispersal barriers. Physically homogenizing cores is therefore not a means of better capturing a single community, it is a mixing of ecologically distinct ones (Ettema 2002; Tecon & Or 2017; Bach *et al*. 2018).

Our results demonstrate that sampling effort and sample placement fundamentally affect diversity estimates. Therefore, maintaining a consistent sampling design is essential in maintaining comparability. However, we also find that true composite samples represent an incomplete portion of the soil microbial community, and the proportion of the community that is represented depends on the spatial heterogeneity of the microbiota, as in most cases, soil cores capture complete communities. This is particularly important when comparing ecosystems, as our results demonstrate that composite sampling can reverse apparent diversity relationships between ecosystems, leading to misleading ecological conclusions. Finally, we find that any additional diversity captured by pooling more cores relative to a single core is significantly undermined when sequencing depth remains fixed regardless of the number of cores pooled. Taken together, our study advocates the sampling and sequencing of individual cores rather than composite samples, especially if the goal is to infer plot-level richness or within-plot heterogeneity. Where compositing is unavoidable, sequencing depth should be scaled to the number of cores composited rather than held at single-core depth. In such cases, (Martiny *et al*. 2006; Chase & Knight 2013; Parnell *et al*. 2025) suggest that pairing composited samples with a subset of individual cores can help retain spatial information, especially in long-term, large-scale monitoring, where unpredictable changes (tree fall, flooding) may force slight changes in the sampling design. These considerations apply equally to transect-based and coordinate-based monitoring designs, where homogenization poses the same fundamental problem regardless of how the sampling unit is defined. Sampling effort and spatial extent determine the scale of inference and fundamentally shape microbial diversity estimates (Chase & Knight 2013; Chase *et al*. 2018), parameters that cannot be adjusted post hoc. Our findings serve to guide the sampling design of soil microbiome assessments, and advocate for the explicit reporting and standardization of sampling effort and spatial extent as fundamental prerequisites for reliable diversity inference.

## Acknowledgments

We thank the managers of the Biodiversity Exploratories, Max Müller, Miriam Teuscher, Franca Marian, and all former managers for their work in maintaining the plot and project infrastructure; Victoria Grießmeier for support through the central office; Andreas Ostrowski for managing the central database; and Markus Fischer, Eduard Linsenmair, Dominik Hessenmöller, Daniel Prati, Ingo Schöning, François Buscot, Ernst-Detlef Schulze, Wolfgang W. Weisser, and the late Elisabeth Kalko for their role in setting up the Biodiversity Exploratories project. We thank the administration of the Hainich National Park, the UNESCO Biosphere Reserve Swabian Alb, and the UNESCO Biosphere Reserve Schorfheide-Chorin, as well as all landowners, for the excellent collaboration. We especially thank Nicole Steinbach for her technical support with the Illumina sequencing workflow and Beatrix Schnabel for her support with sample preparation. This work was funded by the German Research Foundation (DFG) as part of the Priority Programme 1374 “Biodiversity Exploratories” (DFG 512281171)

## Data and Code Availability

The processed sequence data and corresponding metadata are available in the Biodiversity Exploratories Information System BExIS (https://doi.org/10.17616/R32P9Q): sample metadata - BExIS ID 32361, https://doi.org/10.71615/bexis.32361; taxonomic assignments - BExIS ID 32394, https://doi.org/10.71615/bexis.32394; and sequence variant abundance data - BExIS ID 32420, https://doi.org/10.71615/bexis.32420. Raw sequencing data were deposited in the National Center for Biotechnology Information (NCBI) Sequence Read Archive (SRA) under BioProject accession number PRJNA1242586. The R code used to process the data, perform the analyses, and generate the figures is available on GitHub: https://github.com/mikostakou/SpaceMic16S.

## Supplementary materials

**Figure S1.**
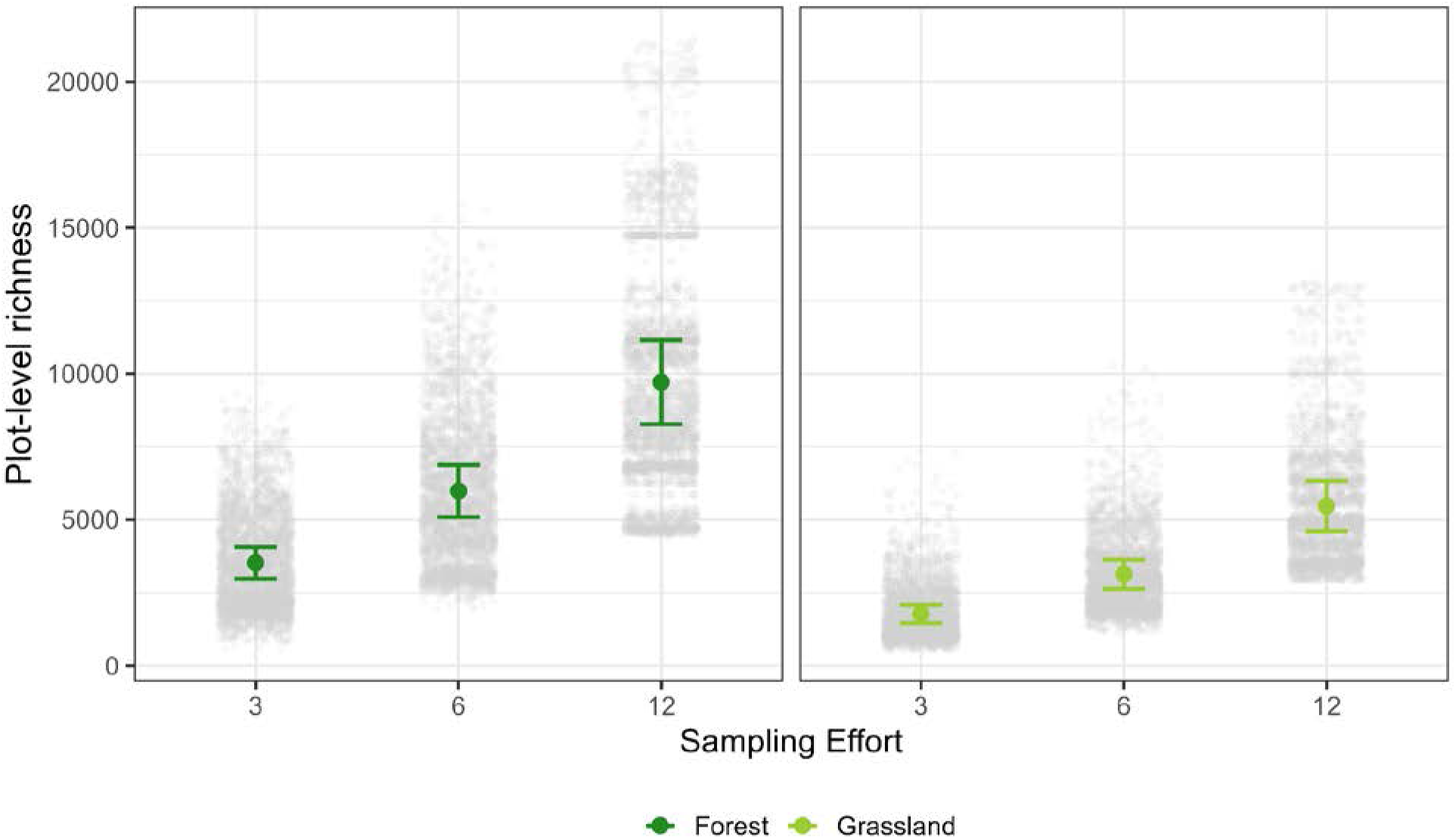
Mean plot-level richness (± SD) in forest and grassland plots. Grey points represent individual sampling configurations, while colored points and error bars show mean plot-level richness values per plot and sampling effort, with 95% confidence intervals calculated across plots.

**Figure S2.**
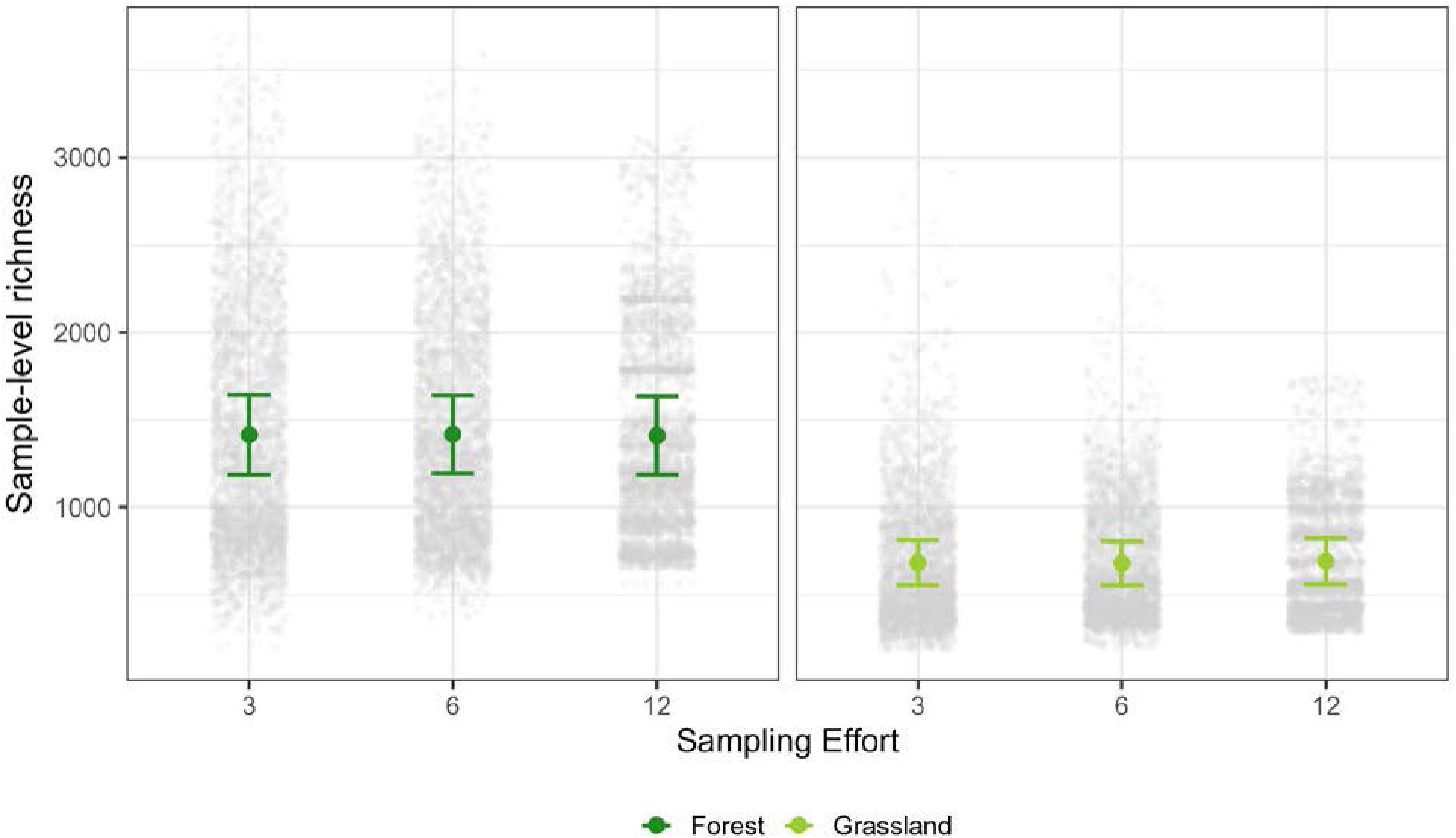
Sample-level richness across sampling efforts (x = 3, 6, 12) in forest and grassland plots. Grey points represent individual sampling configurations, while colored points and error bars show mean sample-level richness values per plot and sampling effort, with 95% confidence intervals calculated across plots.

**Figure S3.**
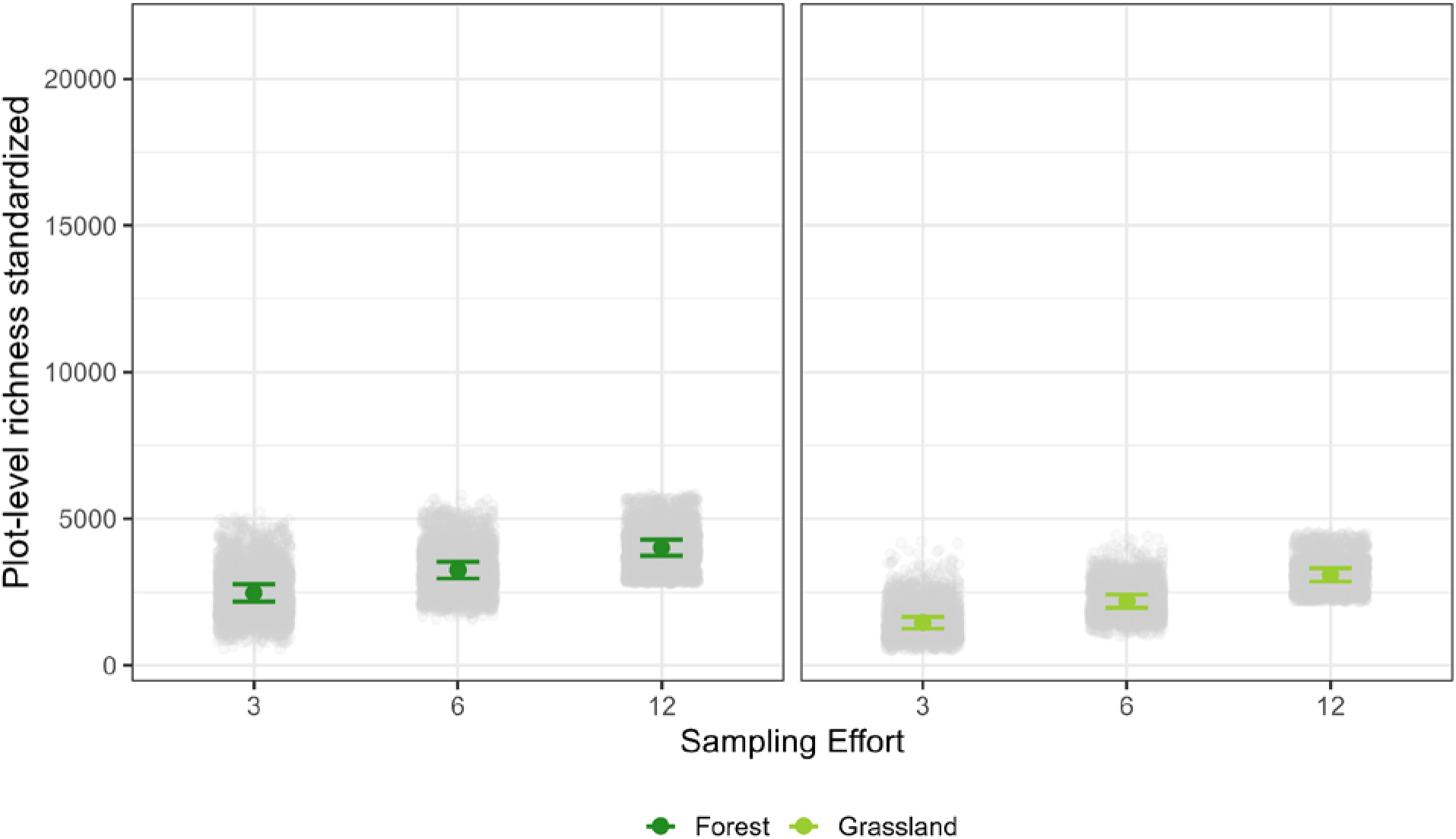
Mean plot-level standardized richness (± SD) in forest and grassland plots. Grey points represent individual sampling configurations, while colored points and error bars show mean values per plot and sampling effort, with 95% confidence intervals calculated across plots.

**Figure S4.**
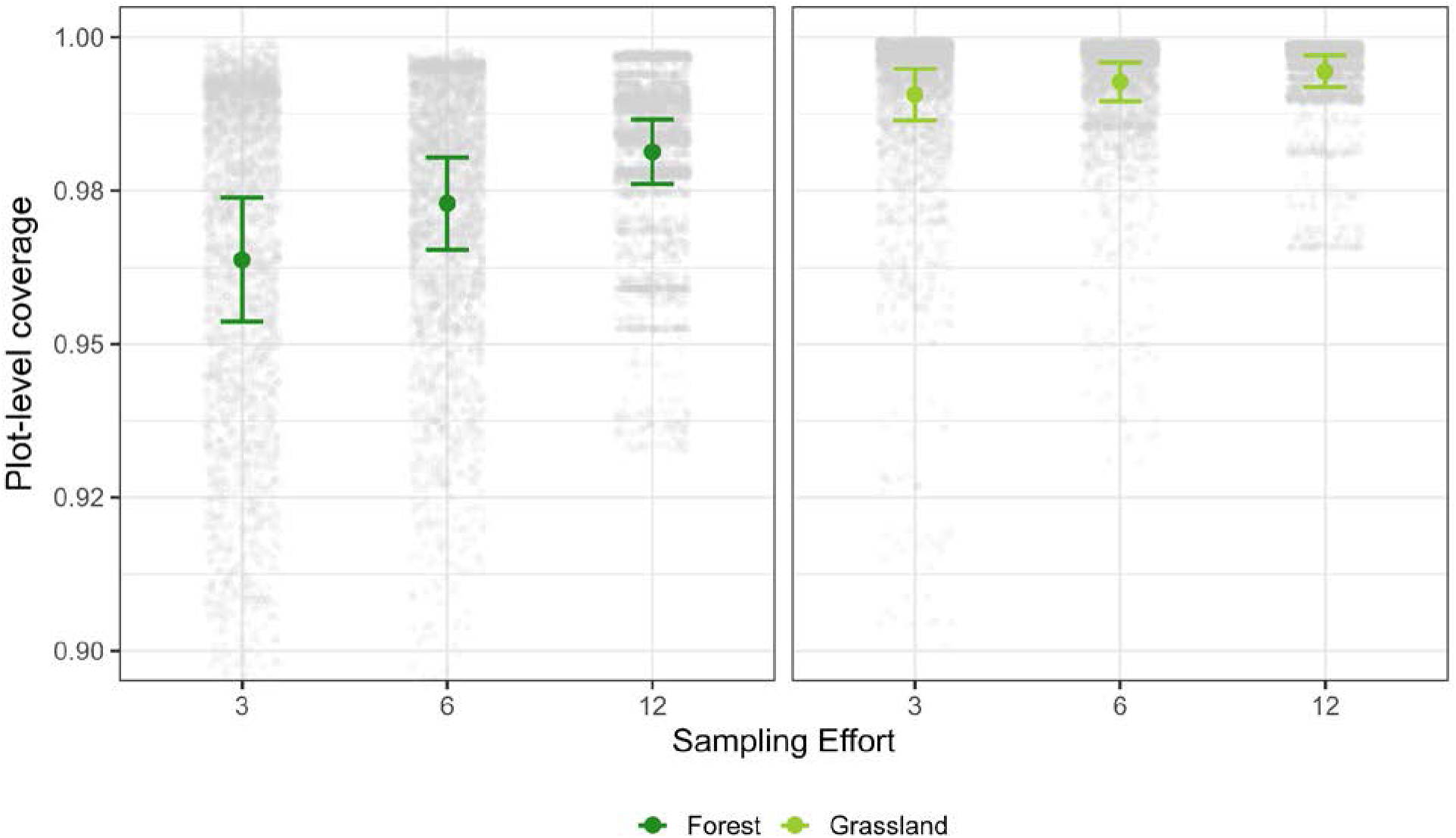
Plot-level coverage across different sampling efforts (x = 3, 6, 12) in forest and grassland plots. Grey points represent individual sampling configurations, while colored points and error bars show mean plot-level coverage per plot and sampling effort, with 95% confidence intervals calculated across plots.

**Figure S5.**
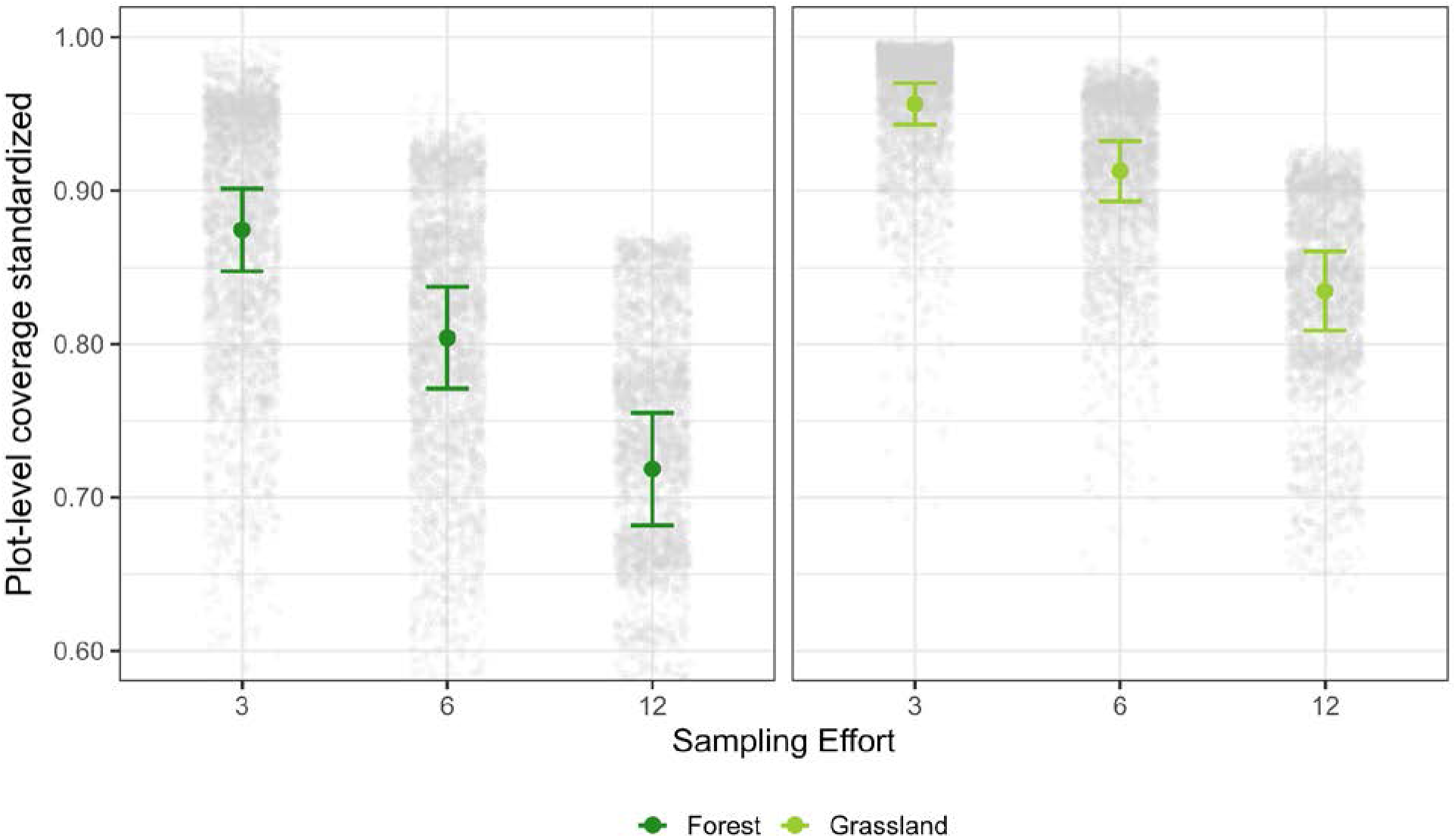
Standardized plot-level coverage across sampling efforts (x = 3, 6, 12) in forest and grassland plots. Grey points represent individual sampling configurations, while colored points and error bars show mean values of standardized plot-level coverage per plot and sampling effort, with 95% confidence intervals calculated across plots.

**Figure S6.**
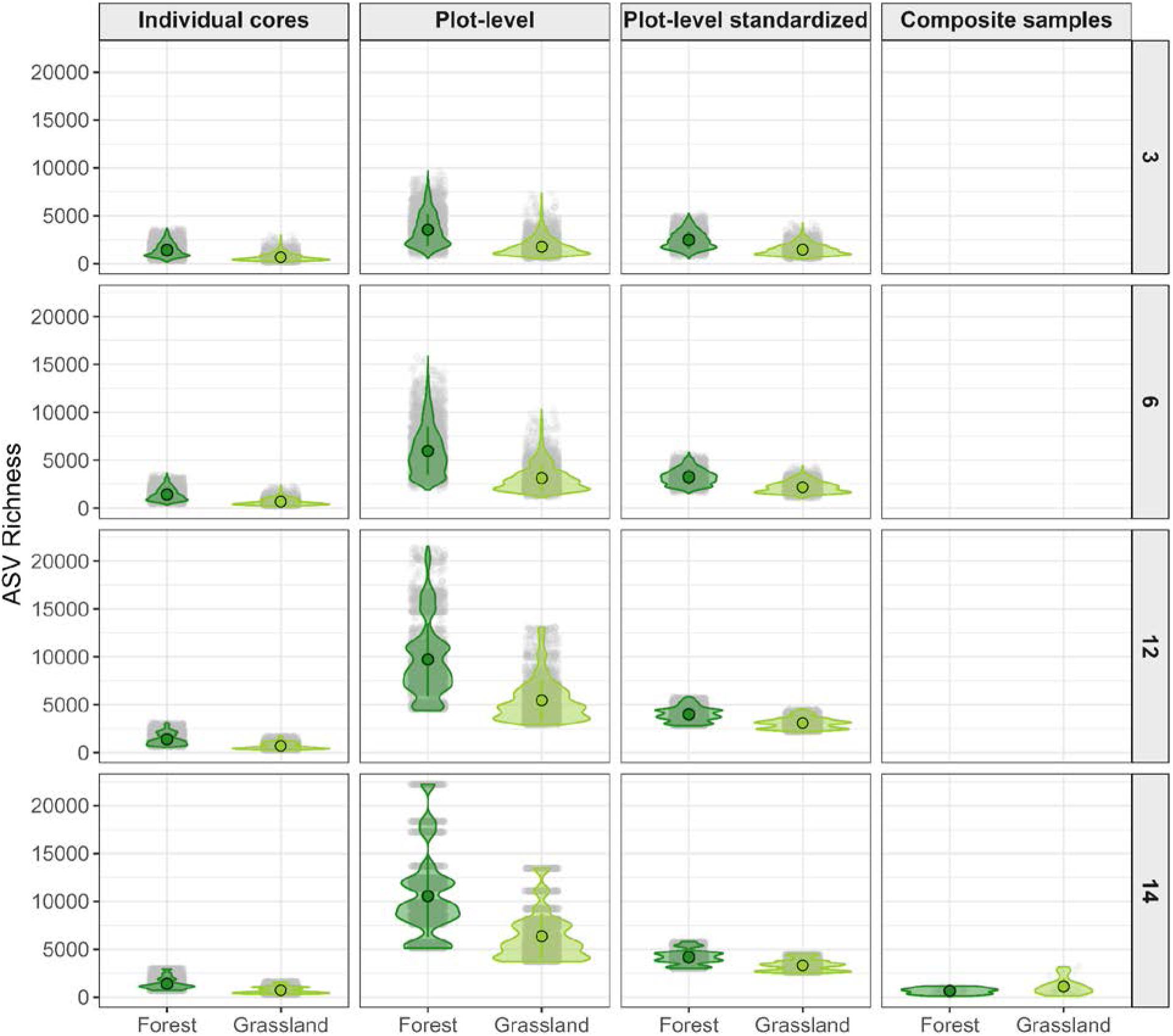
Species richness for selected sampling efforts 3,6,12,14 for forest and grassland plots across sample-level, plot-level, plot-level standardized, and plot-level richness for composite samples.

**Figure S7.**
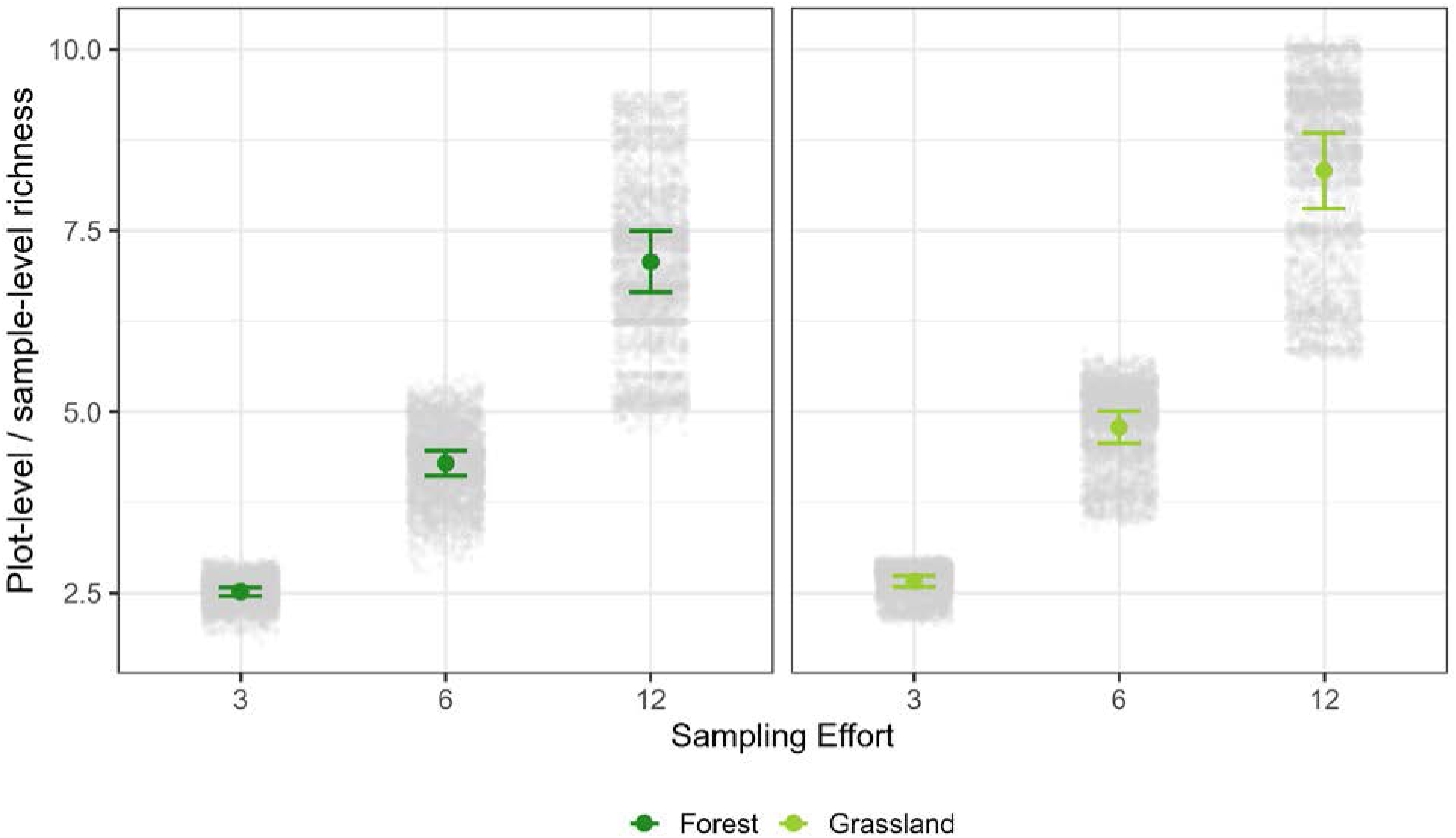
Ratio of sample-level to plot-level richness across sampling efforts (x = 3, 6, 12) in forest and grassland plots. Grey points represent individual sampling configurations, while colored points and error bars show mean values per plot and sampling effort, with 95% confidence intervals calculated across plots.

**Figure S8.**
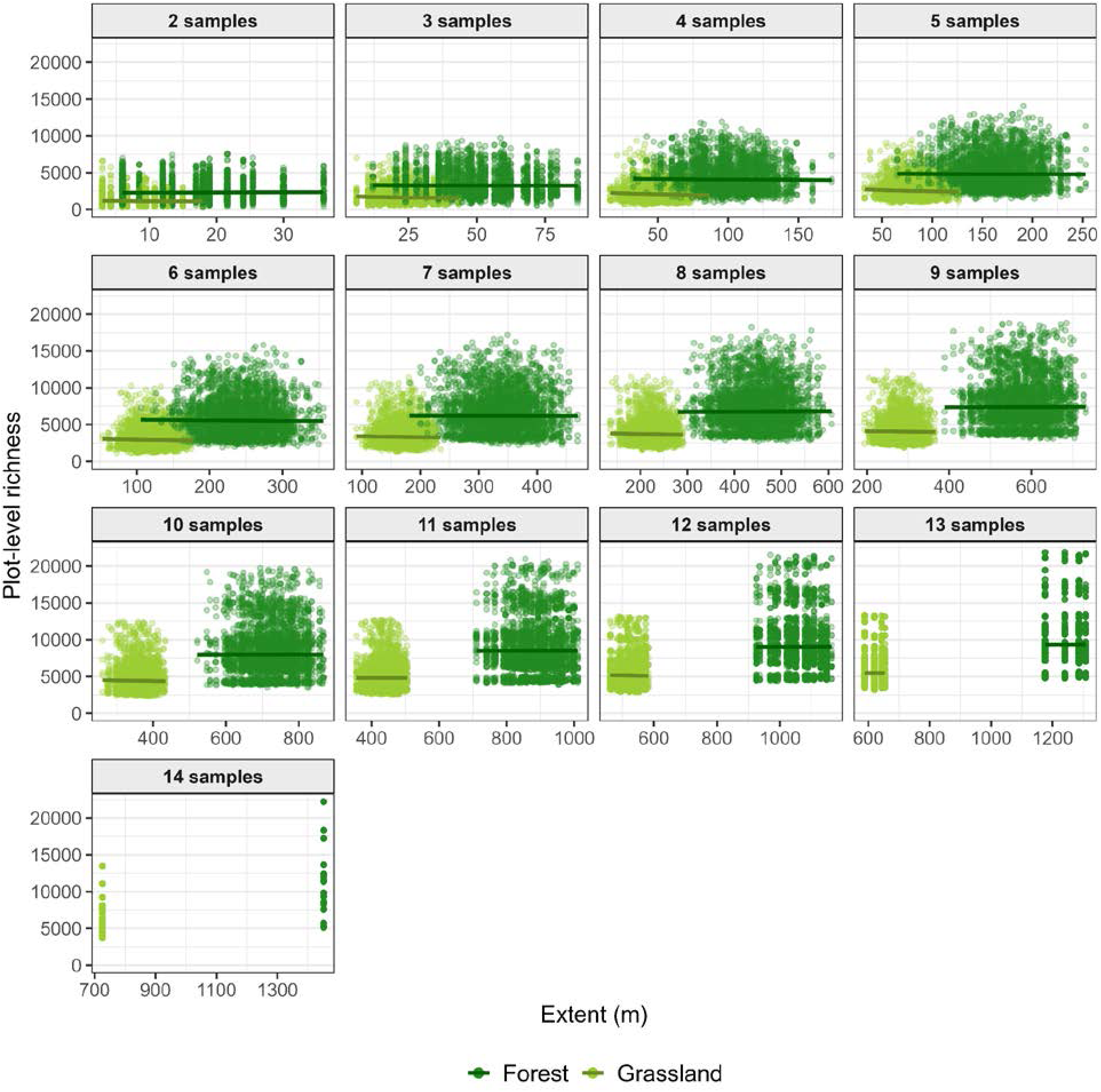
Plot-level richness as a function of spatial sampling extent (m) within the forest–grassland overlap across sampling efforts (2–14 samples). Points show individual sampling configurations, while lines indicate fitted relationships from generalized linear mixed models for each ecosystem type within each sampling effort.

**Figure S9.**
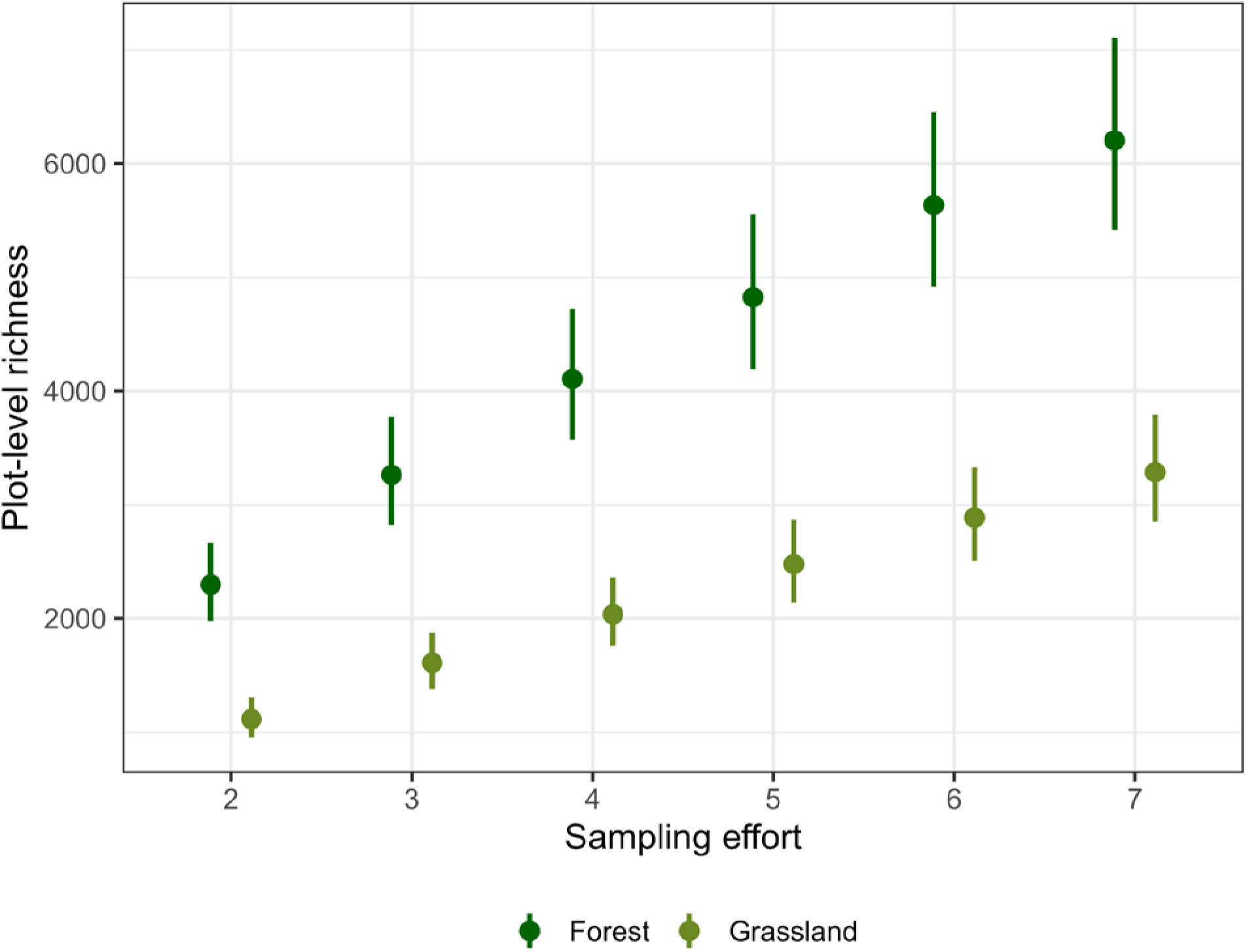
Model-predicted plot-level richness (± 95% CI) for forest and grassland across comparable sampling efforts (x = 2–7). Predictions are derived from generalized linear mixed models fitted separately for each sampling effort.

**Figure S10.**
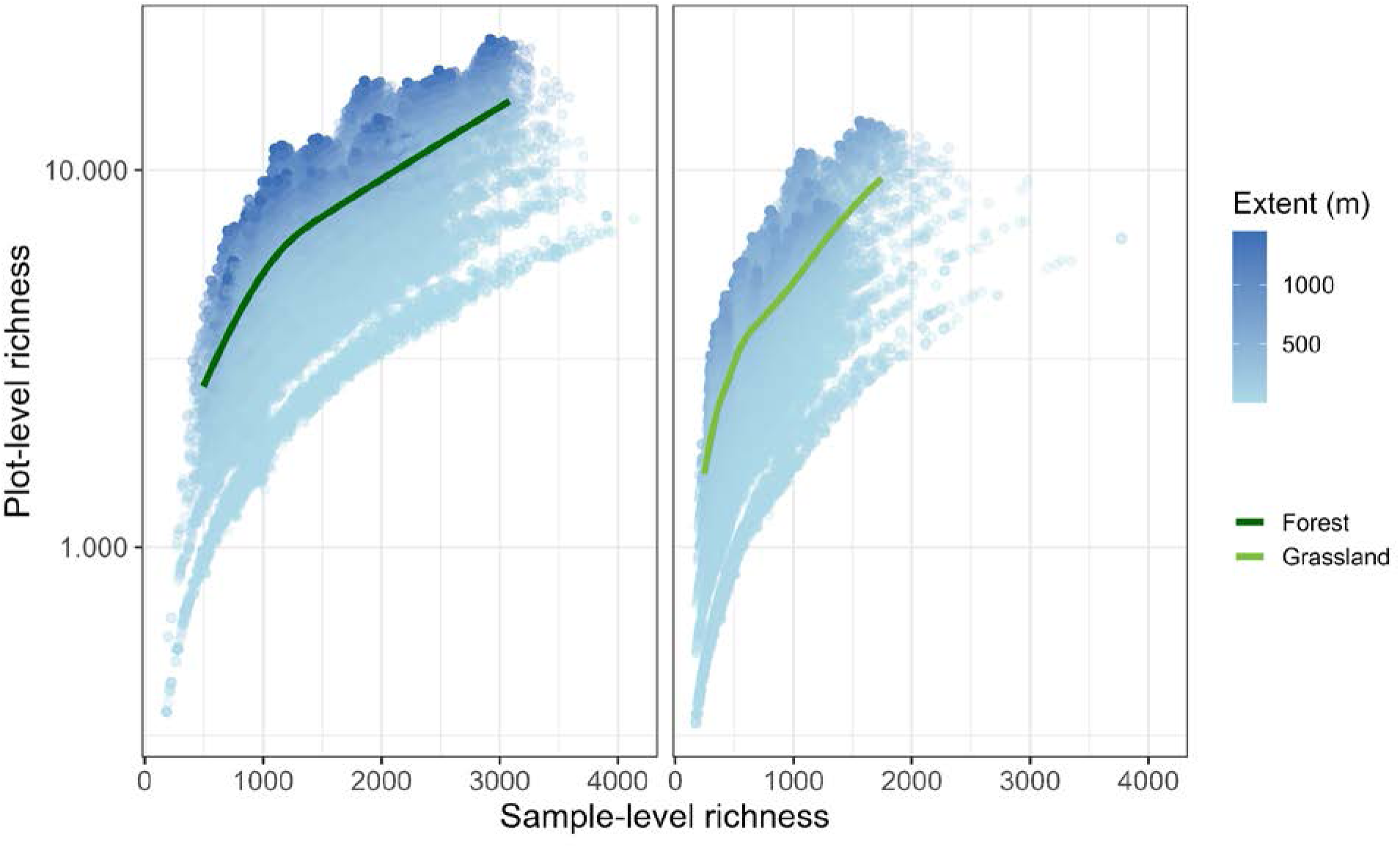
Relationship between sample-level and plot-level richness in forest and grassland plots. Each point represents a single sampling configuration and is colored according to spatial sampling extent (m). Lines represent a generalized additive model (GAM; Gamma family with log link) fit for each ecosystem type. Model predictions are shown at the median sampling extent for each ecosystem.

**Table S1.** Generalized linear mixed model (GLMM) statistics for plot-level richness as a function of sampling effort. Type III Wald χ² tests; plot identity included as a random intercept. Model fit: plot-level richness: AIC = 272,632.8, R²_c_ = 0.931, R²m = 0.627; standardized plot-level richness: AIC = 249,796.8, R²_c_ = 0.890, R²_m_ = 0.579. Significance: *p < 0.05, **p < 0.01, ***p < 0.001.

| Predictor | $\chi^2$ | p-value |
| --- | --- | --- |
| **Plot-level richness** |  |  |
| Intercept | 11,181.634 | < 0.001 *** |
| log(N) | 98,719.813 | < 0.001 *** |
| Ecosystem type | 60.522 | < 0.001 *** |
| log(N) × Ecosystem type | 284.161 | < 0.001 *** |
| **Standardized plot-level richness** |  |  |
| Intercept | 30,021.750 | < 0.001 *** |
| log(N) | 53,810.28 | < 0.001 *** |
| Ecosystem type | 126.45 | < 0.001 *** |
| log(N) × Ecosystem type | 2,138.67 | < 0.001 *** |

x = number of soil samples. Models fitted to sampling efforts 2–14 cores per plot across 57 plots.

**Table S2.** GLMM statistics for plot-level coverage (Good’s Coverage) as a function of sampling effort. Type III W ald χ² tests from Beta GLMMs (logit link); plot identity as random intercept. Model fit: plot-level coverage: AIC = −570,141.8, R²_c_ = 1.000, R²_m_ = 0.431; standardized plot-level coverage: AIC = −297,439.9, R²_c_ = 0.998, R²_m_ = 0.616. Significance: *p < 0.05, **p < 0.01, ***p < 0.001.

| Predictor | $\chi^2$ | p-value |
| --- | --- | --- |
| **Plot-level coverage** |  |  |
| Intercept | 686.925 | < 0.001 *** |
| N (sampling effort) | 15,597.260 | < 0.001 *** |
| Ecosystem type | 57.256 | < 0.001 *** |
| N × Ecosystem type | 1,096.205 | < 0.001 *** |
| **Standardized plot-level coverage** |  |  |
| Intercept | 606.151 | < 0.001 *** |
| N (sampling effort) | 112,111.289 | < 0.001 *** |
| Ecosystem type | 70.527 | < 0.001 *** |
| N × Ecosystem type | 3,621.772 | < 0.001 *** |

x = number of soil cores.

**Table S3.** GLMM statistics for plot-level richness as a function of spatial extent. Fixed-effects coefficient estimates from Gamma GLMMs (log link) with plot identity as a random intercept. Model fit: plot-level richness: AIC = 11,797.2, R²_c_ = 0.928, R²_m_ = 0.622; standardized plot-level richness: AIC = 10,836.2, R²_c_ = 0.885, R²_m_ = 0.590. Significance: *p < 0.05, **p < 0.01, ***p < 0.001.

| Predictor | $\chi^2$ | p-value |
| --- | --- | --- |
| **Plot-level richness** |  |  |
| Intercept | 17,803.536 | < 0.001 *** |
| log(Extent) | 425,146.229 | < 0.001 *** |
| Ecosystem type | 29.193 | < 0.001 *** |
| log(Extent) × Ecosystem type | 798.646 | 0.003 ** |
| **Standardized plot-level richness** |  |  |
| Intercept | 49,686.650 | < 0.001 *** |
| log(Extent) | 242,273.020 | < 0.001 *** |
| Ecosystem type | 101.790 | < 0.001 *** |
| log(Extent) × Ecosystem type | 8,029.340 | < 0.001 *** |

**Table S4.** GLMM statistics for plot-level coverage as a function of spatial extent. Type III Wald χ² tests from Beta GLMMs (logit link) with plot identity as a random intercept. Model fit: plot-level coverage: AIC = −764,263.0, R²_c_ = 1.000, R²_m_ = 0.387; standardized plot-level coverage: AIC = −431,511.9, R²_c_ = 1.000, R²_m_ = 0.638. Significance: *p < 0.05, **p < 0.01, ***p < 0.001.

| Predictor | $\chi^2$ | p-value |
| --- | --- | --- |
| **Plot-level coverage** |  |  |
| Intercept | 992.168 | < 0.001 *** |
| log(Extent) | 11,974.309 | < 0.001 *** |
| Ecosystem type | 73.695 | < 0.001 *** |
| log(Extent) × Ecosystem type | 1,027.916 | < 0.001 *** |
| **Standardized plot-level coverage** |  |  |
| Intercept | 1,723.855 | < 0.001 *** |
| log(Extent) | 128,399.894 | < 0.001 *** |
| Ecosystem type | 83.974 | < 0.001 *** |
| log(Extent) × Ecosystem type | 3,936.230 | < 0.001 *** |

**Table S5.** GLMM statistics for sample to plot-level richness scaling across spatial extent classes. Fixed-effects estimates from Gamma GLMMs (log link), with plot identity as random intercept. Ecosystem type was included as a fixed effect. Model fit: 0–75 m: AIC = 337,881.5, R²_c_ = 0.738, R²_m_ = 0.722, x = 57 plots; 100–300 m: AIC = 267,276.8, R²_c_ = 0.937, R²_m_ = 0.888, x = 57 plots; 600–1,000 m: AIC = 170,995.3, R²_c_ = 0.986, R²_m_ = 0.898, x = 55 plots. Significance: *p < 0.05, **p < 0.01, ***p < 0.001.

| Predictor | Estimate | SE | z | p-value |
| --- | --- | --- | --- | --- |
| **Spatial extent 0–75 m** |  |  |  |  |
| Intercept | −0.065 | 0.063 | −1.03 | 0.301 |
| $\log(\alpha)$ | 1.121 | 0.009 | 121.47 | < 0.001 *** |
| Ecosystem type | 0.016 | 0.061 | 0.27 | 0.789 |
| $\log(\alpha) \times \text{Ecosystem type}$ | −0.029 | 0.009 | −3.25 | 0.001 ** |
| **Spatial extent 100–300 m** |  |  |  |  |
| Intercept | 1.742 | 0.045 | 38.33 | < 0.001 *** |
| $\log(\alpha)$ | 0.970 | 0.006 | 152.28 | < 0.001 *** |
| Ecosystem type | −0.238 | 0.045 | −5.26 | < 0.001 *** |
| $\log(\alpha) \times \text{Ecosystem type}$ | 0.010 | 0.006 | 1.60 | 0.110 |
| **Spatial extent 600–1,000 m** |  |  |  |  |
| Intercept | 2.777 | 0.114 | 24.38 | < 0.001 *** |
| $\log(\alpha)$ | 0.885 | 0.017 | 51.07 | < 0.001 *** |
| Ecosystem type | −0.551 | 0.114 | −4.85 | < 0.001 *** |
| $\log(\alpha) \times \text{Ecosystem type}$ | 0.061 | 0.017 | 3.55 | < 0.001 *** |

$\alpha$ = sample-level richness

**Table S6.** AIC-based model comparison for richness–effort and richness–extent relationships. Models were fitted to plot-level means using Gamma GLMMs (log link) with plot as a random intercept.

| Response variable | Model | AIC | ΔAIC |
| --- | --- | --- | --- |
| **Plot-level richness – sampling effort** |  |  |  |
| Plot-level richness | Log-linear | 11,178.00 | 0.00 |
|  | Asymptotic | 12,153.22 | 975.22 |
|  | Square root | 12,402.22 | 1,224.22 |
|  | Linear | 13,046.17 | 1,868.17 |
| Standardized plot-level richness | Asymptotic | 11,074.36 | 0.00 |
|  | Log-linear | 11,075.19 | 0.84 |
|  | Square root | 11,624.45 | 550.10 |
|  | Linear | 12,023.49 | 949.12 |
| **Plot-level richness – spatial extent** |  |  |  |
| Plot-level richness | Log-linear | 548,830.0 | 0.00 |
|  | Square root | 555,044.4 | 6,214.47 |
|  | Linear | 566,332.9 | 17,502.94 |
|  | Asymptotic | 587,167.1 | 38,337.15 |
| Standardized plot-level richness | Log-linear | 491,700.0 | 0.00 |
|  | Square root | 491,908.6 | 208.62 |
|  | Linear | 508,562.5 | 16,862.58 |
|  | Asymptotic | 521,319.9 | 29,619.90 |

ΔAIC > 2 indicates meaningful support for the better model; ΔAIC > 7 indicates strong support. For standardized richness as a function of sampling effort, the asymptotic and log-linear models were statistically indistinguishable ( ΔAIC = 0.84), consistent with the standardization artefact rather than true biological saturation.

